# Unified AI framework to uncover deep interrelationships between gene expression and Alzheimer’s disease neuropathologies

**DOI:** 10.1101/2020.11.30.404087

**Authors:** Nicasia Beebe-Wang, Safiye Celik, Ethan Weinberger, Pascal Sturmfels, Philip L. De Jager, Sara Mostafavi, Su-In Lee

## Abstract

Deep neural networks offer a promising approach for capturing complex, non-linear relationships among variables. Because they require immense sample sizes, their potential has yet to be fully tapped for understanding complex relationships between gene expression and human phenotypes. Encouragingly, a growing number of diseases are being studied through consortium efforts. Here we introduce a new analysis framework, namely MD-AD (**M**ulti-task **D**eep learning for **A**lzheimer’s **D**isease neuropathology), which leverages an unexpected synergy between deep neural networks and multi-cohort settings. In these settings, true joint analysis can be stymied using conventional statistical methods, which (1) require “harmonized” phenotypes (i.e., measured in a highly consistent manner) and (2) tend to capture cohort-level variations, obscuring the subtler true disease signals. Instead, MD-AD incorporates multiple related phenotypes sparsely measured across cohorts, and learns complex, non-linear interactions between genes and phenotypes not discovered using conventional expression data analysis methods (e.g., component analysis and module detection), enabling the model to capture subtler signals than cohort-level variations. Applied to the largest available collection of brain samples (N=1,758), we demonstrate that MD-AD learns a truly generalizable relationship between gene expression program and AD-related neuropathology. The learned program generalizes in several important ways, including recapitulation of the disease progress in animal models and across tissue types, and we show that such generalizability is not achieved by previous statistical paradigms. Its ability to identify genes with high non-linear relevance to neuropathology enabled us to identify a sex-specific relationship between neuropathology and immune response across microglia, providing a nuanced context for association between inflammatory genes and AD.

## INTRODUCTION

Alzheimer’s disease (AD), the sixth leading cause of death in the United States, is a degenerative brain condition with no known treatment to prevent, cure, or delay its progression. Primary challenges to treating and preventing AD include *extensive heterogeneity* in the clinicopathologic state of older individuals^1^ and *limited knowledge* about genetic and molecular drivers and suppressors of AD-related (amyloid and tau) proteinopathies and AD dementia^2^. Recent efforts to identify molecular mechanisms underlying AD and its progression focus on two complimentary approaches. First, the assembly of large genome-wide association studies (GWAS) (N>100K subjects) enabled case/control analyses of genetic variants correlated with a clinical diagnosis of AD. Interestingly, some identified variants have implicated tau protein binding, amyloid precursor protein (APP) metabolism or immune pathways that play a role in their aggregation and/or uptake^3–5^. These results reinforce the need for detailed investigations of the drivers of neuropathological variation across individuals. Second, moderate-scale post-mortem transcriptomic studies have investigated molecular correlates of a richer set of phenotypic and neuropathological outcomes^6–9^. Early work in this domain examined pairwise correlations among gene expression levels and AD related traits^10^ or a diagnosis of AD^11^. More recent attempts have focused on learning statistical dependencies among gene expression using AD expression data collected from one cohort, in order to infer gene regulatory networks^7^ or co-expressed modules^6^ associated with AD related phenotypes (see Supplementary Methods for details). The relative scarcity of brain gene expression data collected from each cohort has posed a challenge to the use of complex models, such as deep neural networks.

The collection of postmortem brain RNA-sequencing datasets, assembled by the AMP-AD (**A**ccelerating **M**edicines **P**artnership **A**lzheimer’s **D**isease) consortium, provides a unique opportunity to combine multiple data sets in an integrative analysis. Previous work has applied existing co-expression methods to each dataset and used consensus methods to identify consistent gene expression modules across datasets^9^. To our knowledge, there has not yet been a *unified* approach to learn a single joint model that incorporates multiple AMP-AD datasets, which would enable the use of all samples to capture intricate interactions between gene expression levels and phenotypes. A unified approach has been hindered by: (1) the need for “harmonized” phenotypes consistently measured across datasets, and (2) the limitation of current analysis methods that focus on linear relationships between variables (e.g., module analysis^9^) which tend to capture broader patterns in gene expression that often correspond to cohort-level variations, and to consequently obscure true disease signals.

Here, we develop MD-AD (**M**ulti-task **D**eep learning for **A**lzheimer’s **D**isease neuropathology), a *unified framework for analyzing heterogeneous AD datasets* to improve our understanding of expression basis for AD neuropathology (**Figure 1a-d**). Unlike previous approaches, MD-AD learns a single neural network by jointly modeling multiple neuropathological measures of AD (**Figure 1a**), and hence it incorporates a large collection of postmortem brain RNA-sequencing datasets. The combined AMP-AD dataset contains 1,758 samples distributed across 9 brain regions which are labeled with up to six neuropathological outcomes that are *sparsely* available across cohorts (**Figure 1e**). This *unified* framework has key advantages over separately trained models. First, MD-AD can accommodate sparsely labeled data, which is a natural characteristic of datasets aggregated through consortium efforts (**Figure 1e**). Even if different phenotypes only partially overlap in the measured samples, each sample contributes to the training of both phenotype-specific and shared layers (**Figure 1a**). Predicting multiple phenotypes at once biases shared network layers to capture relevant features of these AD phenotypes at the same time. This is of critical importance: each phenotype represents a *different* noisy measurement of the same underlying true biological process, and, as we demonstrate, joint training allows MD-AD to average out the noise to extract the true hidden signal. Additionally, the increased sample size enables MD-AD to capture complex non-linear interactions between genes and phenotypes. Multi-layer perceptrons (MLPs) offer another powerful approach for directly capturing complex relations between gene expression and a phenotype. However, training separate MLPs for each phenotype (**Supplementary Figure 1a**) has limited scope: it can utilize only the samples measured for a specific phenotype, and it cannot share information across related phenotypes. We demonstrate that these advantages improve MD-AD prediction accuracy, enabling its predictions to generalize across species and tissue types (**Figure 1b**).

**Figure 1.**
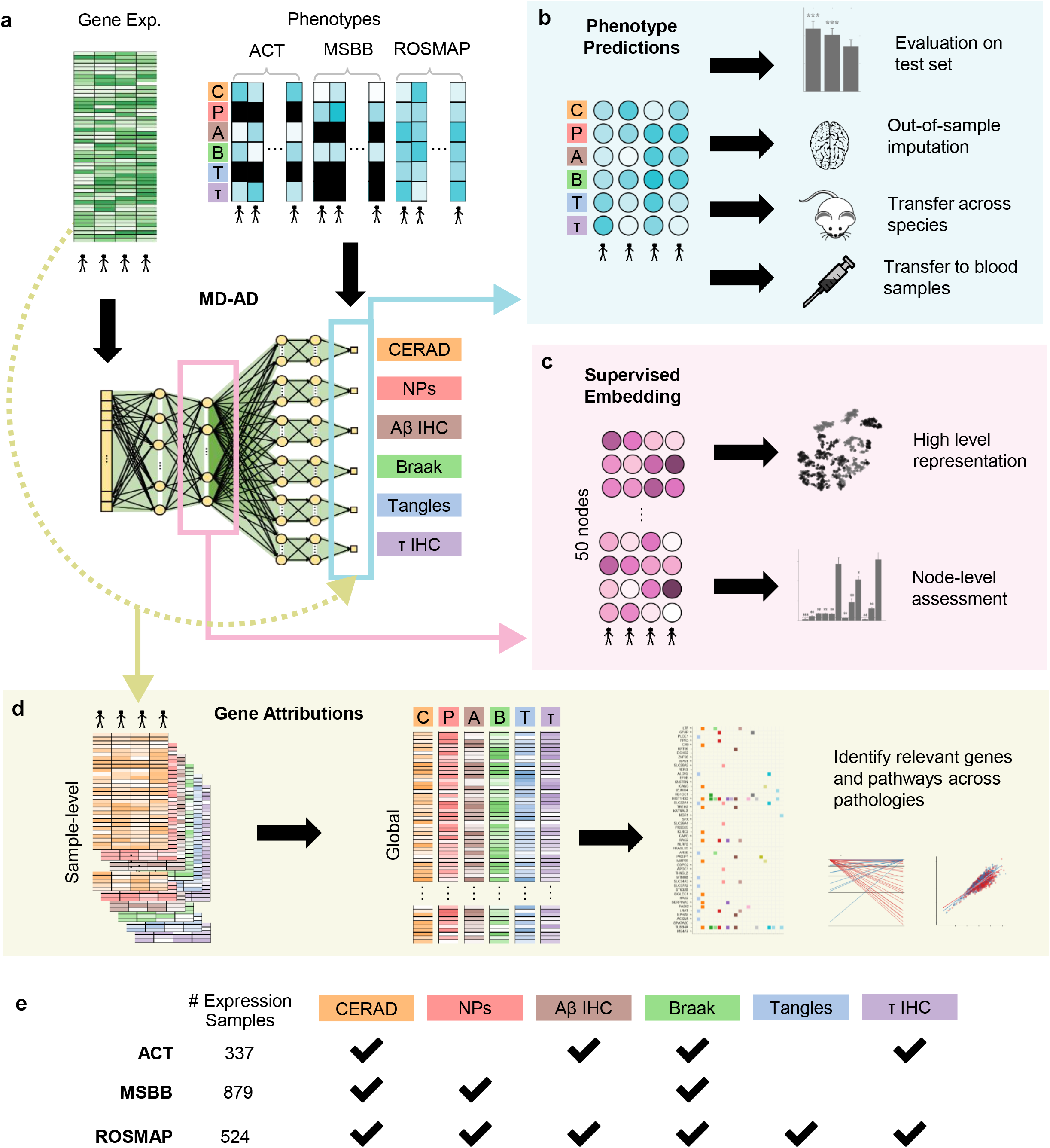
Overview of the MD-AD method and analyses. (**a**) Overview of the MD-AD framework: MD-AD is trained to predict six neuropathology phenotypes simultaneously from brain gene expression samples. During model training, samples do not need to have all available phenotypes; they influence only the layers for which they have labels (including shared layers). (**b**) Illustrates out-of-sample datasets we used to validate MD-AD’s predictions (**c**) Illustrates analyses used to validate the last shared layer of MD-AD. (**d**) By using model interpretability methods, we highlight genes relevant to MD-AD’s predictions. Further analyses reveal non-linear effects among genes and their relationship with AD severity prediction.

MD-AD’s ability to capture complex non-linear relationships provides an opportunity to gain new insights into the expression basis of AD neuropathology, which were not identified by previous approaches. However, an obvious drawback of deep neural networks is their black-box nature, making it difficult to biologically interpret gene-phenotype associations. This paper presents two ways to address this challenge. First, MD-AD adopts a well-known feature attribution method^12^, which quantifies how much each input variable (here, gene expression level) contributes to a prediction (here, a neuropathological phenotype) to identify genes and pathways relevant to each neuropathological phenotype (**Figure 1d**). Second, because MD-AD is a deep learning model, we can interpret its intermediate layers as biologically relevant *high-level feature representation* of gene expression levels and its predictions as the amalgamation of AD-specific molecular markers. The last shared layer of MD-AD can be viewed as *a supervised embedding* influenced by each neuropathological phenotype used during training. Thus, by interpreting this layer’s embedding, we gain understanding of model components and high-level dependencies between expression and neuropathology (**Figure 1c**). As the first deep learning attempt to relate gene expression to multiple AD neuropathological phenotypes, we identify globally important genes not previously implicated in linear methods and perform sex-specific analyses to explore implicitly captured non-linear effects among genes and AD severity predictions.

In sum, our new MD-AD framework makes the following contributions: (1) It is able to *effectively impute accurate AD neuropathological phenotype predictions* from broad compendia of heterogeneous brain gene expression data; (2) it produces learned representations that are more robust than separately learned models, improving *generalizability to other datasets, species, and even tissue types*; (3) it provides an *improved understanding of inter-relationships* among molecular drivers of AD neuropathology that is missed by linear methods; and (4) from a biological standpoint, MD-AD highlights a sex-specific relationship between microglial immune activation and neuropathology.

## RESULTS

### MD-AD provides a unified framework to learn a single model of multiple neuropathological phenotypes across multiple cohort datasets

The MD-AD model takes as input brain gene expression profiles and simultaneously predicts several AD-related neuropathological phenotypes (**Figure 1a**). In particular, the model is trained on expression data from the ROSMAP^6,13,14^, ACT^15^ and MSBB^16^ cohort studies, which together have 1,758 gene expression profiles for 925 distinct individuals. These data are normalized for study batch (Supplementary Methods, **Supplementary Figure 1b-c**)^17^. As shown in **Figure 1a**, the MD-AD model simultaneously predicts six AD-related neuropathological phenotypes: three related to amyloid plaques and three to tau tangles. The former include: (1) **Aβ IHC**: amyloid-β protein density via immunohistochemistry, (2) **NPs:** neuritic amyloid plaque counts from stained slides, and (3) **CERAD score:** a semi-quantitative measure of neuritic plaque severity^18^. The latter include: (4) **τ IHC:** abnormally phosphorylated τ protein density via immunohistochemistry, (5) **tangles:** neurofibrillary tangle counts from silver stained slides, and (6) **Braak stage:** a semi-quantitative measure of neurofibrillary tangle pathology^19^. Thus, MD-AD generates six highly related predictions simultaneously and covers each of the two main hallmarks of AD neuropathology (plaques and tangles) at three levels of granularity. The three studies measure partially overlapping subsets of the six phenotypes described above (**Figure 1e** and **Table 1**), so across our combined dataset some variables are sparsely labeled, although Braak and CERAD are each measured in all studies (**Figure 1e**). During training, the MD-AD model continually updates model parameters via backpropagation, but only for labeled phenotypes from a given sample. Thus, for each phenotype for a given sample, MD-AD updates parameters from associated separate layers along with all shared layers. This allows us to train a unified model from all available samples despite having many missing labels.

### MD-AD accurately predicts neuropathology from gene expression, and its predictions are generalizable to external datasets

In the first pass at model evaluation, we assessed MD-AD using standard five-fold cross-validation (CV), quantifying the average mean squared error (MSE) on the test samples (**Figure 2a and Supplementary Figure 1d**). We compared MD-AD to two simpler baseline models: a regularized linear model (ridge regression) and a single output deep neural network (MLP). These alternative results helped us assess two significant components of the MD-AD model: (1) its non-linear modeling of the relation between gene expression and neuropathological phenotypes, and (2) its joint modeling of multiple related neuropathological phenotypes. In general, MLP models outperformed linear models, highlighting a general advantage of deep learning over a linear approach. Furthermore, compared to the MLP models, MD-AD showed MSE reductions of 7% for CERAD score, 13% for Braak stage, 7% for NPs, 25% for tangles, 10% for Aβ immunohistochemistry (IHC), and 14% for τ IHC (**Figure 2a**). Interestingly, MD-AD showed its largest performance gain for the tangles variable, which also had the most missing labels (**Figure 1e**), highlighting a specific advantage of joint learning for sparsely labeled data.

**Figure 2.**
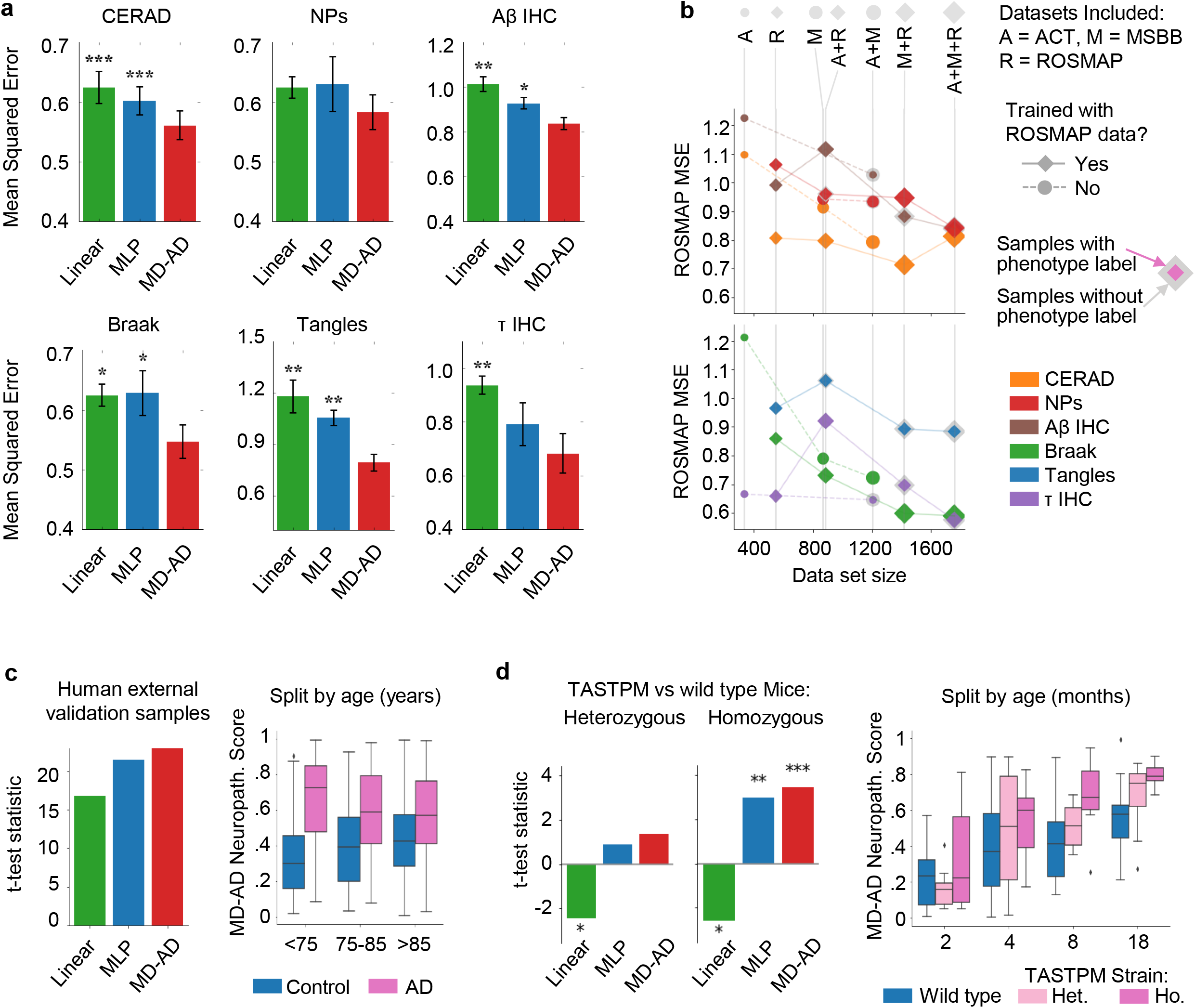
MD-AD prediction performance for within-sample and out-of-sample data. **(a)** Average test set mean squared error (MSE) for phenotype predictions across 5 test splits. MLP: Multiple Layer Perceptron. Linear: linear model using L2 regularization. **(b)** Average MSE for ROSMAP test set samples when training on subsets of the available data sets in the training set. **(c)** For samples from three external validation data sets, we obtain neuropathology scores for each sample by averaging the percentiles of predictions across all six neuropathology variables. *Left*: t-test statistics measuring the difference between each model’s predicted neuropathology scores for AD-diagnosed vs. control individuals. All tests results were statistically significant (*p*<.001). *Right*: Box plots displaying the distribution of MD-AD’s predicted neuropathology scores split by age group and diagnosis (see **Supplementary Figure 3b** for sample sizes and significance of pair-wise differences). **(d)** *Left*: t-test statistics measuring the difference between each model’s predicted neuropathology score for heterozygous TASTPM vs. wild type mice. *Middle*: t-test statistics measuring the difference between each model’s predicted neuropathology score for homozygous TASTPM vs. wild type mice (*: *p*<.05, **: *p*<.01, ***:*p*<.001). *Right*: Box plots displaying the distribution of MD-AD’s predicted neuropathology scores for mice split by age and strain (See **Supplementary Figure 3c** for sample sizes and significance of pair-wise differences).

Because our model was trained and evaluated on ACT, MSBB, and ROSMAP datasets, we assessed whether residual (uncorrected) batch effects affected performance. To do so, we performed additional validation experiments by leaving out specific datasets during training and then evaluating their performance for MD-AD trained on the other datasets (**Figure 2b, Supplementary Figure 2a**). We evaluated MSE performance for ROSMAP alone since it was the only dataset with all six phenotype labels; further, by evaluating a single dataset’s performance, we can identify the influence of adding “external” data. We make several observations from this analysis. First, as one may expect, larger training samples always helped reducing prediction error on test samples from the unseen study (ROSMAP), and especially so when datasets from multiple cohorts were included in the training (i.e., ACT and MSBB) (circular markers in **Figure 2b**). Second, when considering the effects of augmenting ROSMAP data with other datasets during training (diamond markers in **Figure 2b**), we observed that errors initially increased when adding a new dataset but tended to decline as more datasets were included in training. This may result from small differences in labeling conventions across studies, or batch effects in gene expression data. However, we find that the benefits of additional heterogeneous samples ultimately outweigh potential batch effects in prediction performance. Third, interestingly, we observed that adding new samples improved performance for a phenotype even when the phenotype in question was not measured in the new samples (see gray footprints around markers in **Figure 2b**). This suggests that the shared representation learned by MD-AD captures the underlying biological signal common across noisy neuropathological phenotype measurements.

Next, as the ultimate test of MD-AD out-of-sample predictions, we assessed performance on three independent studies never seen by the model: Mount Sinai Brain Bank Microarray (MSBB; N=1,O53), Harvard Brain Tissue Resource Center (HBTRC; N=460), and Mayo Clinic Brain Bank (N=323). Because these datasets provide a sparse set of neuropathological labels, we evaluated whether MD-AD predictions were consistent with the (binary) neuropathological diagnosis of AD by calculating “MD-AD neuropathology scores” for each sample (by averaging ranked predictions across the six phenotypes). For comparison with other methods, we also generated “neuropathology score” predictions for our baseline models.

As shown in **Figure 2c**, we observed a highly significant difference in predicted neuropathology scores between AD cases and controls (two-sided t-test: *t*= 22.98, *p*<0.001), and these differences were more pronounced for MD-AD compared to the other baseline models (results split by dataset are shown in **Supplementary Figure 3a**). More convincingly, when split by age group (**Figure 2c** right panel), we consistently observed a significant increase in predicted neuropathology for AD vs control samples, but the difference was largest in individuals under 75 (between-groups *p*-values are shown in **Supplementary Figure 3b**). This is consistent with the observation that aging individuals who are cognitively nonimpaired often have substantial neuropathology^15^. Together, these results indicate that MD-AD can identify generalizable gene expression patterns that are predictive of AD-related neuropathology across varied age ranges, and thus it is unlikely that these patterns merely capture normal aging.

### Complex transcriptomic predictors of neuropathology are conserved across species

We next evaluated how well MD-AD’s learned expression patterns predictive of neuropathology recapitulated neuropathology in mouse models. We applied MD-AD trained on human datasets to make predictions based on brain (hippocampal and cortical) gene expression data from 30 TASTPM mice that harbored double transgenic mutation in APP and PSEN1 and compared the predictions to those from 76 wild type mice^20^. We focused on TASTPM mice because they were found to robustly exhibit early signs of amyloid aggregation and plaque formation. As above, to simplify MD-AD predictions, we then predicted all six neuropathological phenotypes via MD-AD and generated an aggregate “neuropathology score” per mouse (as described in Supplementary Methods).

As shown in **Figure 2d**, MD-AD predicted significantly higher neuropathology scores for the homozygous cross TASTPM than wild type mice (two-sided t-test: *t*=3.45, *p* <.001). The MLP baseline method also produced significant differences between homozygous and wild type mice, but less effectively (*t*=3.01, *p*<.01). Furthermore, there was a stronger trend for higher predictions in the heterozygous TASTPM cross (N=32) than wild type mice for MD-AD (*t*=1.38, *p*=.17) compared to MLP baselines (*p*=.38). Interestingly, our linear baseline tended to predict lower average neuropathology levels for these AD strains than wild type, suggesting that a linear approach may fail to effectively model crossspecies AD signal. None of the models produced significantly different neuropathology scores between other strains (i.e., TPM, TAS10, Tau) and wild type mice, consistent with lower neuropathological burden in these models (data not shown). Notably, when we stratified the samples by age, we found that MD-AD tended to predict higher neuropathology in older mice regardless of strain), but in particular it made higher neuropathology predictions for homozygous than heterozygous crosses followed by wild type mice (many of these groups differed significantly from one another, as shown in **Supplementary Figure 3c**). Overall, these results indicate that MD-AD learns a generalizable expression pattern associated with neuropathology that is conserved across species.

### Deep transcriptomic signatures of neuropathology are predictive of AD dementia

Hidden layers of a deep neural network capture the embedding of input examples in the derived feature space, yielding a “hidden” representation that is predictive of the outcome(s) of interest. In this case, the last shared layer of MD-AD (**Figure 1a, c**) captures a latent (lower) dimensional representation of gene expression that is predictive of multiple types of neuropathology related to AD. To derive the biological basis of MD-AD predictions, we first visualized this embedding space in 2D using the t-SNE algorithm (**Figure 3a**)^21^ (to improve stability, we used a consensus approach over many re-trainings of the MD-AD model, **Supplementary Figure 4a**). We observed that the representation in this space was impressively coherent with respect to all six neuropathological variables: individuals with similar overall neuropathology severities had similar MD-AD consensus representations for their gene expression profiles, and this observation was true for external test samples not used for model training (**Figure 3d-e, Supplementary Figure 3d**). This was remarkable because representations derived by unsupervised dimensionality reduction (e.g., K-means or PCA) failed to capture the components of gene expression relevant to neuropathology, and mainly captured batch effects, while those derived by standard single output MLP tended to overfit to each neuropathology variable and were incoherent *across* neuropathological measurements (**Figure 3c** and **Supplementary Figure 5**).

**Figure 3.**
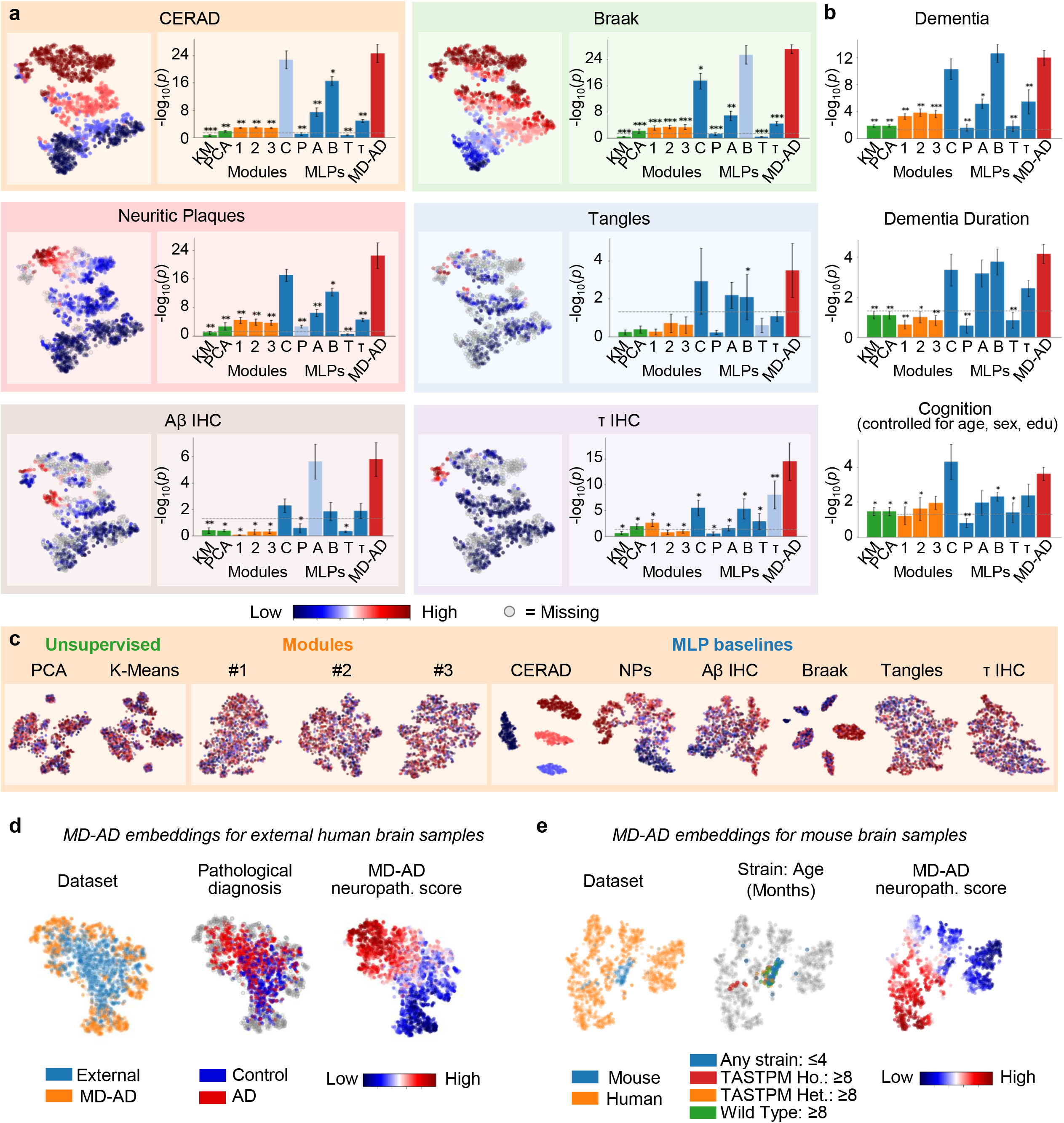
Comparing MD-AD’s supervised embedding to other embedding methods. **(a)** For each colored box, *Left:* 2-dimensional t-SNE embedding of MD-AD’s last shared layer colored by neuropathological phenotype indicated in the title of the box, *Right:* −log_10_(*p*-value) of correlations between “best” node from each embedding method and the neuropathological phenotype across 5 test folds. The “best” node was identified as the most significantly correlated in the training set, but the figure reports correlation - log_10_(*p*-value)’s in their corresponding test sets. Bar graph columns (left to right): two unsupervised embeddings (green; K-Means and PCA), three module-based embeddings (orange; Modules #1^7^ and Modules #2^6^, and Modules #3^9^), six singly-trained MLPs (blue), and MD-AD (red). **(b)** Highest correlation −log_10_(*p*-values) (averaged across 5 training folds) found between each embedding method and high-level AD phenotypes: dementia (diagnosis prior to death), dementia duration (approximate time between dementia diagnosis and death; available for ACT and ROSMAP), and last available cognition score (controlling for age, sex and education; available for ROSMAP only). All *p*-values listed are shown after FDR correction over the nodes within each method. **(c)** 2-dimensional t-SNE embedding of alternative embedding methods (described in **a**). **(d)** 2-dimensional t-SNE embeddings of MD-AD embeddings for training and external data sets. Each point represents a sample colored by dataset (Left), AD status for external samples (Middle) and MD-AD’s predicted neuropathology score (Right). (**e**) 2-dimensional t-SNE embeddings of MD-AD embeddings for external human and mouse samples.

Next, we evaluated whether the MD-AD embedding can go beyond neuropathology to also capture the molecular manifestation of AD dementia. In particular, we considered three “higher-level” phenotype variables: AD dementia (a *clinical* diagnosis of AD), assessment of cognitive function, and assessment of AD duration. We then correlated the latent representation captured by the hidden nodes in the last shared layer with each of these three higher-level phenotypes. As shown in **Figure 3b**, we found that MD-AD consistently produced nodes that were significantly correlated with high-level AD phenotypes; using paired *t*-tests, these correlations often outperformed nodes from our MLPs and always outperformed unsupervised methods and module-based approaches (*p*<.05 after FDR correction over nodes). This indicates that MD-AD creates embeddings that most consistently capture the relationship between gene expression and general AD severity. Together, these results show that by jointly predicting several neuropathological phenotypes, the MD-AD framework produces a low dimensional representation of gene expression data, in the form of embedding nodes, that robustly captures a generalizable signature of AD beyond individual neuropathological phenotypes alone. Detailed annotations for MD-AD embedding nodes are provided in **Supplementary Table 2** and **Supplementary Figure 4b-d**.

### MD-AD reveals an interrelationship between sex and immune genes predictive of AD neuropathology

We next sought to interpret MD-AD’s learned parameters to identify the set of genes (and their relationships) that underlie its impressive predictive performance. Here, we applied the Integrated Gradient (IG) algorithm^12^ on the fully trained model in an ensemble fashion to ensure robustness (Supplementary Methods, **Supplementary Figure 6a-b**), producing an “importance score” for each gene. For a global view, we first performed functional enrichment analysis (GSEA^22,23^) using these importance scores, and found that relevant genes for the MD-AD model were enriched for several pathways, including metabolism of RNA and proteins, immune system, cell-to-cell communication, and signal transduction (**Figure 4b**). **Figure 4a** shows the top 50 genes and their pathway annotations where the particular relevance of immune function is even more prominent.

**Figure 4.**
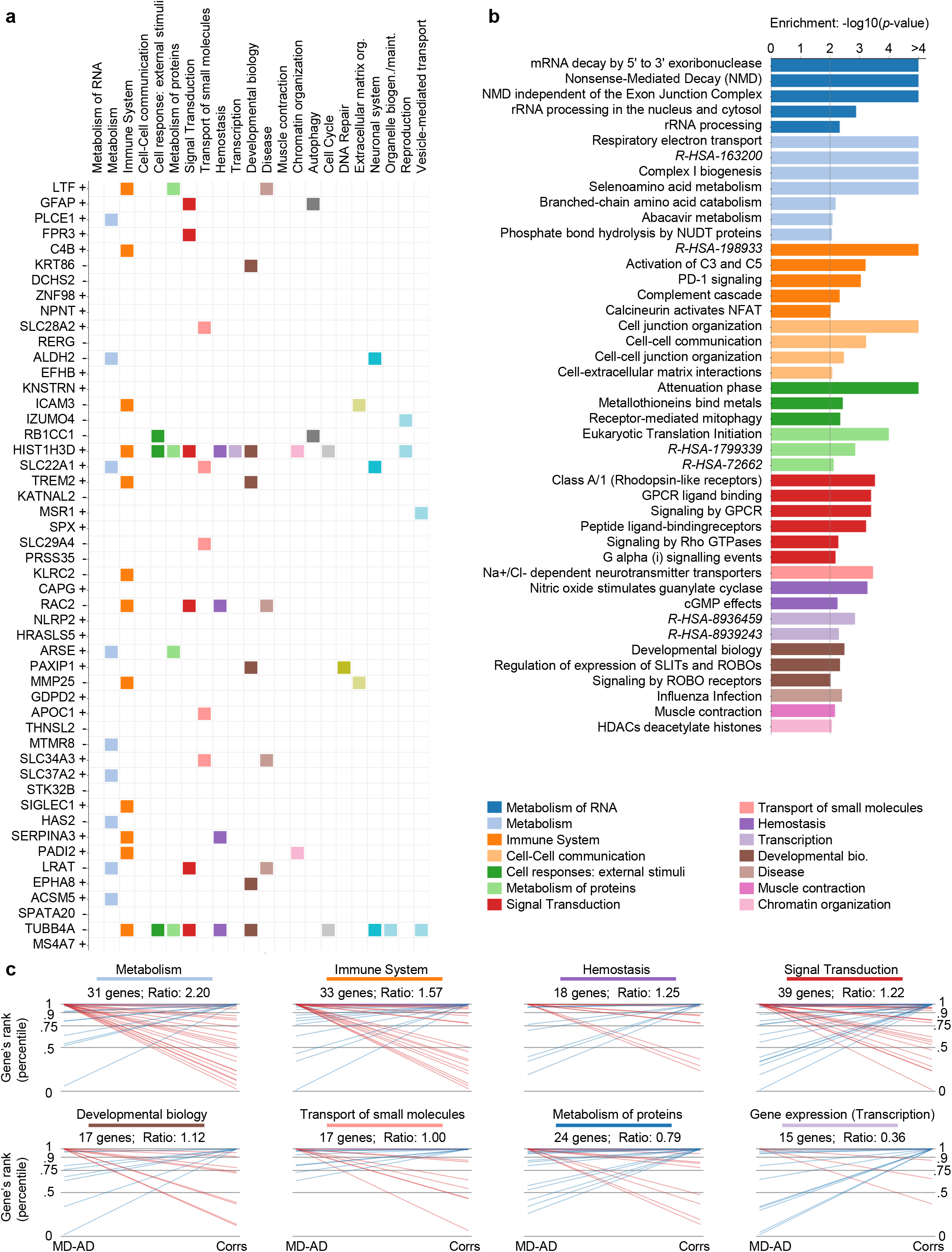
Top predictive genes for the consensus MD-AD model. **(a)** Top 50 MD-AD genes and whether they are negatively (-) or positively (+) associated with high neuropathology. Colored squares indicate that the gene belongs at least one pathway in the column-labeled REACTOME category. **(b)** Gene set enrichment −log_10_(*p*-value) across the final MD-AD gene ranking for REACTOME pathways. Bars are colored by the pathway’s REACTOME category. We show all pathways with significant enrichment (*p*<.01). REACTOME pathways with long names are indicated by their REACTOME stable IDs. **(c)** Comparison of top genes from MD-AD vs a linear correlation-based approach. For each ranking method, we identify the top 1% of all genes and check their membership in REACTOME categories. For each REACTOME category with at least 15 genes in the top 1% of MD-AD and/or correlation rankings, we generate the following plot: each line represents a gene, with left endpoint at the percentile rank for MD-AD and right endpoint at percentile rank for correlations. For clarity, we color the line purple if the gene falls in the top 1% of both MD-AD and correlations, red if it is only in the top 1% of MD-AD, and blue if it is only in the top 1% of correlations. Finally, the title indicates the ratio of MD-AD to correlation-based top genes for the given REACTOME category.

We next assessed to what extent the learned gene importance varied between a linear model and a non-linear model like MD-AD. With a simple linear correlation-based gene ranking, we found that the top 50 genes had a much lower prevalence of REACTOME pathways (**Supplementary Figure 7a**). When we directly compared the top 1% of genes from MD-AD versus a correlation-based approach in **Figure 4c**, we observed that many genes belonging to metabolism, immune system, and signal transduction pathways were highly ranked for MD-AD but not for correlation-ranking. In contrast, transcription-related genes were more frequently highly ranked for correlation-based rankings compared to MD-AD’s rankings. Overall, gene importance scores generated via correlations alone were enriched for a much larger set of REACTOME categories (**Supplementary Figure 7b**), whereas MD-AD pathways tended to be more specific (**Figure 5b**). We saw similar results when performing the same analyses with KEGG pathways (**Supplementary Figure 8**)^24^.

**Figure 5.**
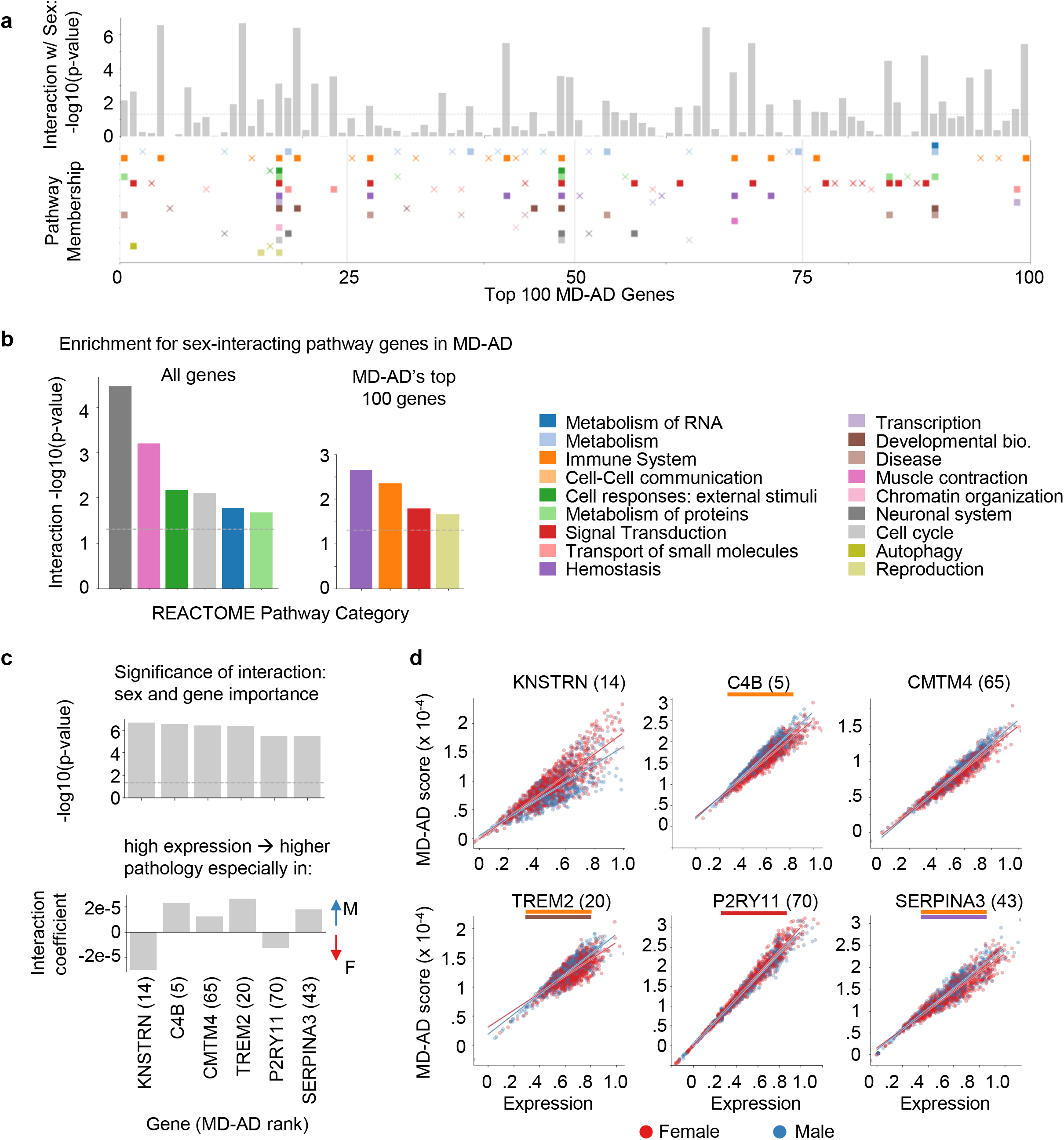
MD-AD’s top genes and their interactions with sex. **(a)** For the top 100 MD-AD genes, we compute the significance of the interaction between expression and sex for its MD-AD score. The bars indicate the gene’s −log_10_(p-value) of the interaction term with sex (after FDR correction), and pathway categories each gene belongs to are indicated below. A filled square indicates that the gene significantly interacts with sex (*p*<.05 after FDR correction), and an “x” marker indicates that it does not. **(b)** For genes with significant sex interactions, we compute the significance of the overlap between REACTOME category genes and sex-differential genes among: Left: all genes, and Right: the top 100 MD-AD genes only. **(c)** For the top 100 MD-AD genes, we identify the genes with the most significant sex interaction for MD-AD scores. We show the significance of the interaction (Top) and the interaction coefficients (Bottom) for the top 6 most sex-differential genes. Each gene’s MD-AD rank is indicated in their x-axis labels **(d)** For the top 6 most-sex differential top 100 MD-AD genes, we display scatter plots of expression by MD-AD score, coloring each sample by sex of the donor.

The nonlinear relationships identified by MD-AD can implicitly capture interaction effects with other covariates observable from expression data (e.g., sex, age, medication intake). Leveraging the fact that, if our model captures a nonlinear effect, then two samples with the same expression level for a single gene could receive different IG (“importance”) scores by MD-AD (e.g., **Figure 5d**; in contrast, a linear model would have no vertical dispersion), we assessed whether a covariate like sex could explain discrepancy between expression levels and IG scores. (Sex is a major risk factor in AD and has prominent gene expression signatures^25^). Thus, we modeled each gene’s IG score as a linear combination of the gene’s expression, the individual’s sex, and the interaction between them to identify sex-interacting genes relevant to AD. Of the 14,591 genes in our dataset, 6,465 showed differential MD-AD importance between sexes (*p*<0.05 after FDR), demonstrating that sex-specific expression effects in AD may be widespread. To confirm that genes are not sex-differential by chance, we show the distribution of sex-differential genes compared with the same analysis conducted with shuffled sex labels (**Supplementary Figure 9a**). However, we were particularly interested in genes with high overall MD-AD importance. When focusing on the top 100 genes with the highest MD-AD scores, we consistently observed high degrees of interaction between sex and immune system genes (as well as reproduction and hemostasis-related genes) (**Figure 5a-b**; we saw similar patterns for KEGG pathways in **Supplementary Figure 9b-c**).

We next explored specific examples of genes with high MD-AD rankings and strong interactions with sex (i.e., the six genes from the top 100 MD-AD list with the strongest interaction *p*-values; **Figure 5c-d**): *KNSTRN, C4B, CMTM4, TREM2, P2RY11*, and *SERPINA3*. In particular, for each of these genes, we observed high expression values associated with higher neuropathology predictions but some stratification across sexes: high expression in females led to especially high neuropathology predictions for *KNSTRN* and *P2RY11*, while the opposite was true for the other four genes. More broadly, our finding immune genes display sex-differential contributions to MD-AD scores appears to be consistent with conclusions from recent studies about sex differences in neuroinflammatory activity and the role these differences may play in neurodegenerative disorders^26^.

We note that some of our top sex-interacting genes may play important roles in immune response, particularly in microglia. *TREM2*, which is genetically implicated in AD, interacts with *CD33* (another AD susceptibility gene)^27^, is an important contributor in the clearance of toxic Amyloid-β by microglia in mice^28^, and is correlated with Aβ deposition in the human brain^27^. Similarly, *KNSTRN* is known to be upregulated in mouse microglial cells’ early response to neurodegeneration^29^. These findings indicate that MD-AD may capture patterns related to sex-differential microglia activity. To explore this idea further, we obtain lists of upregulated genes from nine clusters of single cell microglial transcriptomes^30^, and compare them to our MD-AD gene rankings. As expected, many top MD-AD genes are upregulated in multiple microglial clusters (**Figure 6a**); correlation-based methods ranked these microglial genes less highly (**Supplementary Figure 9d**). Furthermore, genes upregulated in clusters related to stress, immune function and proliferation tended to be sex-differential in their gene importance (**Figure 6b**), further strengthening the finding that sex differences in immune response and inflammation may be an important factor in the molecular basis of age-related neuropathology.

**Figure 6.**
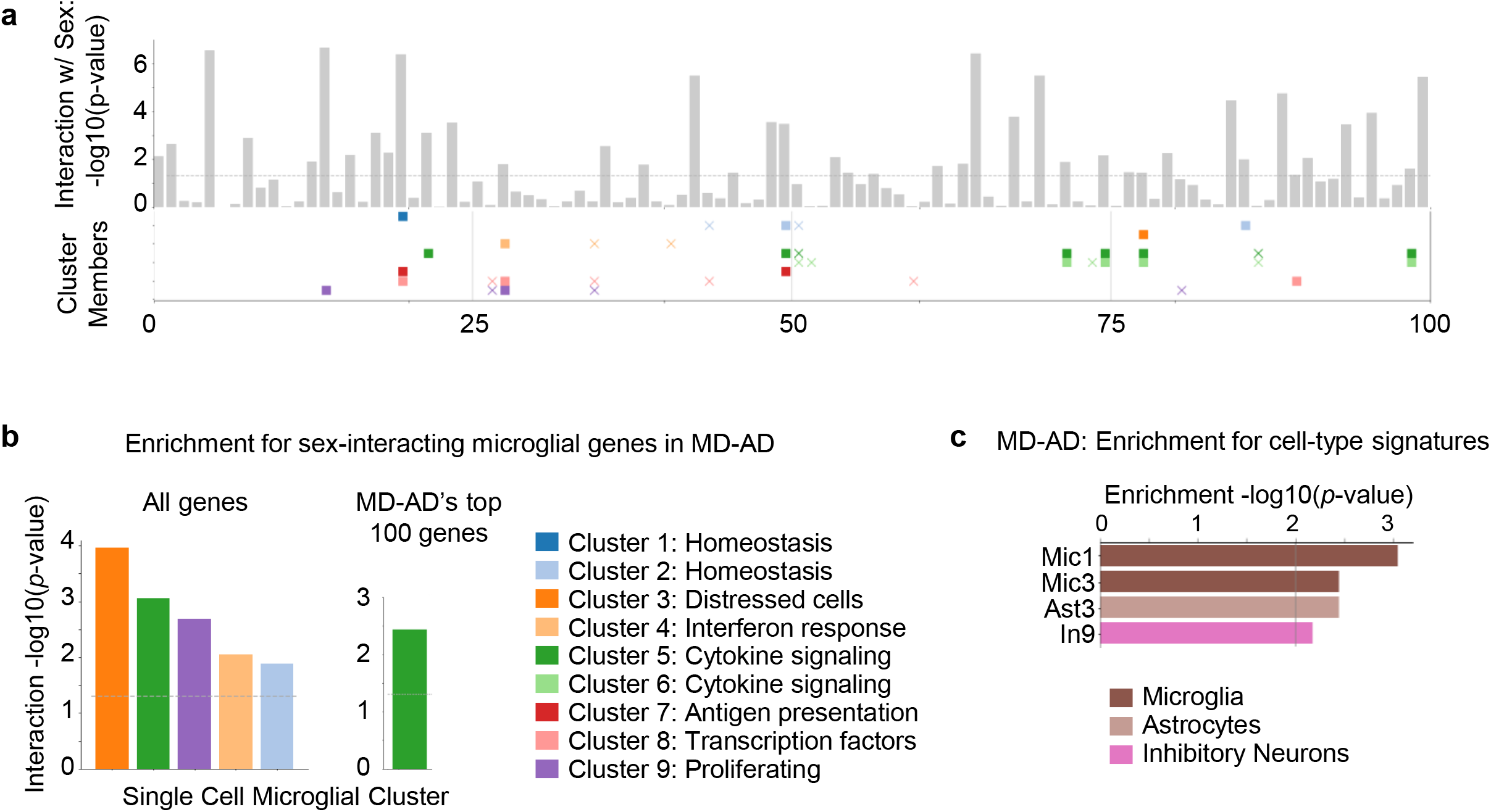
MD-AD’s reliance on microglial cluster genes and gene set signatures. **(a)** Bars indicate the gene’s −log_10_(p-value) of the interaction term with sex (after FDR correction), and gene membership in microglial cluster gene sets from Olah et al.^30^ is indicated below. A filled square indicates that the gene significantly interacts with sex (*p*<.05 after FDR correction), and an “x” marker indicates that it does not. **(b)** For genes with significant sex interactions, we compute the significance of the overlap between microglial cluster genes and sex-differential genes among: Left: all genes, and Right: the top 100 MD-AD genes only. **(c)** Gene set enrichment −log_10_(*p*-value) across the final MD-AD gene ranking for cell type signatures.^8^

To more broadly identify possible cell-type specific effects of MD-AD’s important genes, we tested for the enrichment of 41 different cell type clusters (across six cell types) found by a single cell transcriptomic analysis of AD^8^. Here, we found an enrichment of 2 different microglia clusters, as well as astrocytes and inhibitory neuron clusters (**Figure 6c**). Hence, MD-AD’s predictions of neuropathology rely on broader transcriptomic events that goes beyond microglia genes, suggesting a heterogeneity in the underlying molecular biology that is predictive of accumulation of AD-related neuropathology.

### Complex transcriptomic predictors learned by MD-AD are conserved across tissues

Although MD-AD was developed for brain gene expression data, we next asked whether the learned transcriptomic signatures generalize to blood. To this end, we applied our brain-trained MD-AD model to gene expression datasets from two batches of the AddNeuroMed cohort, which we called Blood1 and Blood2 (NCBI GEO database accessions GSE63060 and GSE63061, respectively; summarized in **Supplementary Table 3**)^31^. As shown in **Figure 7a**, MD-AD predicted significantly higher neuropathology scores for individuals with both mild cognitive impairment (MCI) (two-sided t-test: *t*=7.34, *p* <.001) and AD dementia (two-sided t-test: *t*=5.87, *p* <.001) compared to cognitively normal controls (CTL). Consistent with external brain samples shown in **Figure 2d** and **2f**, MD-AD predictions tended to increase with age for cognitively normal individuals, while they were consistently significantly higher for MCI and AD individuals compared to controls for individuals under 80 years old (**Figure 7b, Supplementary Figure 10b**). Importantly, we noted that a linear model failed to make meaningful predictions (**Figure 7a** and **Supplementary Figure 10a**), suggesting that complex models like MD-AD have better performance in extracting the true underlying signal transferrable between tissues than linear models.

**Figure 7.**
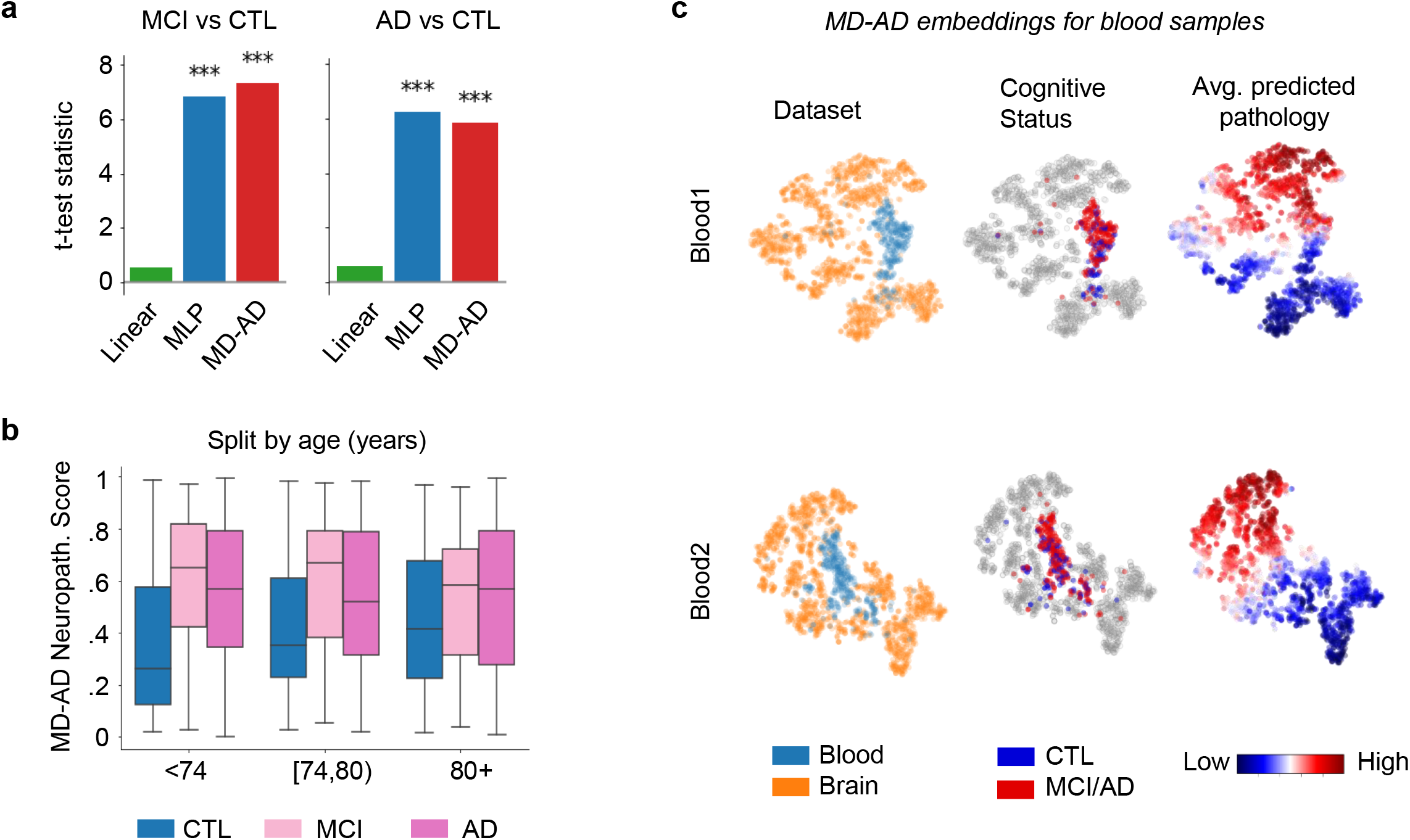
MD-AD’s transfer performance for blood gene expression data sets. **(a)** Shows t-test statistics comparing average predicted neuropathology between individuals with mild cognitive impairment (MCI; Left) and Alzheimer’s dementia (AD; Right) vs. cognitively normal (CTL) individuals. **(b)** Box plots show the differences in predicted neuropathology for blood samples from individuals stratified by age group and cognitive status. Significant differences are shown in **Supplementary Figure 10b. (c)** t-SNE embedding of last shared layer from MD-AD models trained for Blood1 and Blood2 datasets. Samples are colored by their dataset (Left), cognitive status (while brain samples are shown in grey; Middle), and predicted neuropathology score (Right).

Next, we evaluated whether the patterns captured by the MD-AD model were consistent across training brain gene expression samples and blood. To this end, we again visualized MD-AD’s learned embedding using the t-SNE algorithm (**Figure 7c**). We noted a clear difference in expression patterns between blood and brain samples (as seen by the clustering of blood samples in **Figure 7c**); however, MD-AD nevertheless produced an embedding for blood data that stratified blood samples along predicted neuropathological phenotypes in a manner highly consistent with the blood donor’s cognitive status (**Figure 7c; Supplementary Figure 10c**). Together, these analyses indicate that jointly learning the relationship among brain gene expression and several neuropathological phenotypes may allow for learned representations that span tissues. This in turn can open up new avenues for early identification of individuals at risk, and provide new clues into tissue-agnostic molecular mechanisms underlying AD dementia.

## DISCUSSION

We introduce MD-AD, a deep neural network approach for jointly modeling the relationship between brain gene expression data and multiple sparsely labeled neuropathological phenotypes in a multi-cohort setting. By exploiting the synergy between deep learning and a multi-cohort, multi-task setting, we demonstrated that MD-AD can capture complex, non-linear feature representations that are not learned using conventional expression data analysis methods. Specifically, we observed that multi-task learning improves prediction performance over singly trained models. Adding data from different cohorts improves performance for various phenotypes, even those that lacked labels. When we extended our method to other datasets, it captured AD-related biological signals, showing that MD-AD can transfer effectively to out-of-sample, out-of-species (mouse), and even out-of-tissue (blood) datasets.

As a neural network framework, MD-AD’s last shared layer embedding reveals high-level features of gene expression that are predictive of neuropathology according to the intermediate components of the model. As expected, due to multi-task supervision, our embedding nodes tend to relate to AD-associated neuropathology far more effectively than do standard unsupervised approaches and earlier reported (unsupervised) module-based approaches. Compared to singly task-supervised neural networks, the joint training MD-AD performs consistently provided a more stable and coherent AD-related embedding. By exploring the molecular pathways relevant to each node, we identified relevant gene sets contributing to these high-level AD-related features of gene expression.

Finally, we leveraged the complex relationships learned by MD-AD to refine our understanding of the molecular drivers of AD neuropathology. By interpreting genes relevant to our model’s predictions, we uncovered that MD-AD relied on many genes not found in earlier linear-based methods, including several immune system genes. These findings expand the general narrative established by human genetic studies of AD and now a proteomic study of AD^32^; in particular, we see enrichment for complement pathway genes (**Figure 4**) which likely connect with the role of the complement receptor 1 *(CR1)* gene which harbors an AD susceptibility variant whose functional consequences remain poorly understood but do include an influence on the accumulation of neuritic plaque pathology^33–36^. Thus, MD-AD results converge with human genetic results to emphasize the role of complement in AD; interestingly complement protein C4B emerges as one of the top pathology-related genes that display a strong interaction with sex, with men showing a much stronger association than women (**Figure 5c**). This is similar to the behavior of TREM2, another well-validated AD susceptibility gene (**Figure 5c**); however, its relation to amyloid pathology in ROSMAP data was previously reported as being modest^27^. MD-AD was able to uncover its more prominent role in transcriptional data, which is obscured by its sex-dependent nature. Likewise, women reported to have higher expression of a signature of aged microglia in these data^26^, and two modules of co-expressed cortical genes enriched for microglial genes and associated with amyloid (module m114) or tau (module m5) pathology are also influenced by sex^37^. However, the role of neither group of genes is explained by sex; this indicates that the role of sex in the impact of the immune system in AD is complex. MD-AD was able to uncover this complexity more effectively, as is illustrated in **Figure 5c** where some genes have greater effects in men and other in women. Thus, it is not the case that role of the immune system is polarized in one of the two sexes; rather, some pathways and perhaps certain cell subsets may have a larger role in women while others are dysfunctional in men. This could explain why the role of immune genes is more prominent in our analyses: reports from simpler linear models often included immune pathways^6^ but other pathways usually figured more prominently in these earlier RNA-based network models. A meta-analysis of RNA studies (which include the ROSMAP data) highlighted the larger number of sex-influenced genes among the AD-associated gene modules and noted that microglial cells appear to be enriched for both male and female-specific expression effects. With our list of results and our careful evaluation of sex effects we now have an important new road map with which to guide our exploration of the role of microglia in AD in a sex-informed manner. This perspective will be critical not only for mechanistic studies whose results could be obscured by sex effects but also, more importantly, by guiding the study design of clinical trials as highly targeted therapeutic agents emerge to modulate the immune system in AD.

This is but one of the narratives that has emerged from our initial deployment of the MD-AD approach in the aging brain. As new cohorts are characterized, sample sizes expand and new data such as single nucleus RNA sequencing profiles emerge, our approach will help to facilitate data integration and to uncover insights that would not otherwise emerge. Beyond enabling good predictions, our report may actually highlight a more important contribution of MD-AD in resolving key elements of the data structure in the nodes that we defined: these are more than simple aggregates of factors with predictive power. They are beginning to uncover complex interactions, such as the impact of sex which is involved in both men and women, but in different ways, making it difficult to appreciate the role of certain immune pathways in simpler statistical models.

## SUPPLEMENTARY METHODS

### 1. DATA PROCESSING

For developing the MD-AD model, we used data from the following RNA-Seq and neuropathology datasets available through the AMP-AD Knowledge Portal: (1) Adult Changes in Thought (ACT)^15^, (2) Mount Sinai Brain Bank (MSBB)^16^, and (3) Religious Orders Study/Memory and Aging Project (ROSMAP)^6,13,14^. Details of sample collection and sequencing methods are described in previously published work^6,13–16^. We pooled together brain gene expression data from the temporal cortex, parietal cortex, hippocampus, and forebrain white matter from ACT, Brodmann areas 10, 22, 36, and 40 from MSBB, and the dorsolateral prefrontal cortex from ROSMAP. To avoid confounding conditions, we excluded samples from individuals who had neuropathological diagnoses other than AD. Taken together, the studies provide 1,758 gene expression samples.

In order to compile gene expression samples across the three cohorts, we retain expression levels for genes which are present in all datasets. Within each dataset, we exclude genes with null values for over two-thirds of samples. Before combining datasets, we log-transformed the expression values and then normalized them for each gene to vary between 0 and 1. We then combined the gene expression datasets and performed batch effect correction with ComBat^17^ to reduce systematic differences across studies (**Supplementary Figure 1b-c**)^17^. The resulting dataset contains 1,758 gene expression samples, each with 14,591 genes measured.

Next, for each gene expression sample, we incorporated the available corresponding neuropathology labels: (1) **Aβ IHC**: amyloid-β protein density via immunohistochemistry, (2) **plaques:** neuritic amyloid plaque counts from stained slides, and (3) **CERAD score:** a semi-quantitative measure of neuritic plaque severity^38^, (4) **τ IHC:** abnormally phosphorylated τ protein density via immunohistochemistry, (5) **tangles:** neurofibrillary tangle counts from silver stained slides, and (6) **Braak stage:** a semi-quantitative measure of neurofibrillary tangle pathology^19^. Detailed descriptions for each phenotype within each dataset are provided in **Supplementary Table 1**. Because Braak stage and CERAD score are global measurements of neuropathological damage, if an individual had multiple available gene expression measurements from different regions, they each sample was labeled with the same Braak and CERAD values. However, Aβ-IHC and τ-IHC were provided for several brain regions for both ROSMAP and ACT studies. Therefore, each expression sample was labeled with the Aβ-IHC and τ-IHC measurements for the same or nearest region. Because the available plaques label provided by MSBB was averaged over several brain regions, we similarly used ROSMAP’s average plaques and tangles labels (aggregated from several regions) for consistency with MSBB’s metrics (see **Supplementary Table 1**). Finally, for consistency across datasets, we first normalized all neuropathological variables to vary between 0 and 1 before combing datasets.

### 2. COMPUTATIONAL METHODS

#### A. Review of previous approaches

Post-mortem transcriptomic studies have investigated molecular phenotypic and neuropathological outcomes in AD. Early work in this domain examined simple correlations among gene expression and AD symptoms^10^ or compared gene expression levels across AD-patients versus controls^11^. More recently, more systematic network-based analyses have contributed to the understanding of AD biology. In particular, Zhang et al.^7^ constructed molecular networks based on bulk gene expression data separately for individuals with and without AD, and identified modules with remodeling effects in the AD network. More recently, Mostafavi et al.^6^ used co-expressed genes in the aging human frontal cortex to build a single molecular network and identified modules related to AD neuropathological and cognitive endophenotypes. Using single-cell RNA sequencing data, Mathys et al.^8^ clustered cells within brain cell-types to identify and characterize AD-related cellular sub-populations. Each of these approaches have been applied to single cohorts. Until recently, a unified and robust modeling of AD neuropathology based on brain gene expression has been hindered by relative scarcity and regional heterogeneity of brain gene expression datasets. One possible solution is to combine multiple data sets to gain statistical power. The collection of postmortem brain RNA-sequencing datasets, assembled by the AMP-AD (**A**ccelerating **M**edicines **P**artnership **A**lzheimer’s **D**isease) consortium, provides new opportunities to combine multiple data sets. However, such heterogeneous datasets pose challenges to many methods, which must account for inter-study differences. In a recent attempt, Logsdon et al.^9^ used a meta-analysis approach to identify co-expressed modules separately for 7 brain regions across 3 datasets, then subsequently applied consensus methods to identify modules that were conserved across multiple regions and studies. As of now, we’re not aware of any methods that directly model all data in a unified way.

#### B. The MD-AD Model

MD-AD (**M**ulti-task **D**eep learning for **A**lzheimer’s **D**isease neuropathology), is a *unified framework for analyzing heterogeneous AD datasets* to improve our understanding of expression basis for AD neuropathology (**Figure 1**). Unlike previous approaches, MD-AD learns a single neural network by jointly modeling multiple neuropathological measures of AD severity phenotypes, and hence can incorporate data collected from multiple datasets. This *unified* framework has key advantages over separately trained models. First, MD-AD allows sparsely labeled data, which is a natural characteristic of datasets aggregated through consortium efforts (**Figure 1e**). Even if different phenotypes only partially overlap in the measured samples, each sample contributes to the training of both phenotype-specific and shared layers. Predicting multiple phenotypes at once biases shared network layers to capture relevant features of these AD phenotypes at the same time. This is of critical importance: each phenotype represents a *different type* of noisy measurement of the same underlying true biological process, and as we demonstrate by joint training MD-AD is able to average out the noise to extract the true hidden signal. Additionally, the increased sample size enables MD-AD to capture complex non-linear interactions between genes and phenotypes. In contrast, Multi-layer perceptrons (MLPs) offer another powerful approach for directly capturing complex relations between gene expression and a phenotype. However, training separate MLPs for each phenotype (**Supplementary Figure 1a**) has limited scope: it can utilize only the samples measured for a specific phenotype, and it cannot share information across related phenotypes. We demonstrate that these advantages improve MD-AD prediction accuracy, enabling it predictions to generalize across species and tissue types (**Figure 1b**). As illustrated in **Figure 1a**, the MD-AD network jointly predicts six neuropathological phenotypes from gene expression input data via shared hidden layers followed by task-specific hidden layers.

### 3. TRAINING & EVALUATING MD-AD

As described above, we build the MD-AD model in Python using the TensorFlow and Keras packages. In order to have efficient and robust training and to reduce overfitting, we apply a principal component analysis (PCA) transformation to the data and use resulting top 500 principal components – a 500-dimensional representation of our 14,591 gene expression values – as the input to the MD-AD and all baseline models. For comparison to MD-AD, we generate six analogous MLP networks with un-shared representations, and six linear models containing no hidden layers, to serve as baseline models (see **Supplementary Figure 1a**).

In order to robustly evaluate the performance of the models, we segment the dataset into five parts, and each part is treated as a test set once. Within each of the five training and test split splits, each model architecture was trained and hyperparameter-tuned using five-fold cross validation within the training set. We then train each model with the best hyperparamters found by cross validation using the full training set before performance was evaluated on the corresponding test set (see **Supplementary Figure 1d**). Thus, prediction performance reported in the results section are the average of these five test performance values. For training the models, we use a mean squared error (MSE) loss function applied to each phenotype prediction. For the MLP and linear baselines, parameters of the networks are updated via back-propagation for 200 epochs from the mean-squared error (MSE) of the network’s prediction on the given variable’s label among training batches. Similarly, MD-AD’s parameters are also updated via back-propagation, with the loss function calculated as the sum over MSEs across all six prediction tasks (masking losses for missing phenotypes). For MD-AD, we explored several different options for architectures with different amounts of shared and task-specific layers (**Supplementary Figure 2b-c**). We selected the final architecture (shown in **Figure 1a**) because we wanted to have multiple hidden layers in both the shared portion and task-specific portion of the network to allow for non-linear interactions to be learned in both the shared representation and in the task-specific branches, and **Supplementary Figure 2b-c** shows that alternatives to this approach tended to perform similarly or worse.

#### A. Internal test-set validation

As described above, for each training and test split, we use five-fold cross-validation to make modeling choices for the MD-AD model and baselines before training each model with the full training set and reporting and reporting test MSEs (averaged over all five test splits). We evaluate model performance in two ways: (**1**) standard train and test sets, and (**2**) ROSMAP test performance for different subsets of the available datasets.

First, separately for each of our five cross validation training sets, we calculate the final test MSE on the corresponding hold-out set. To test whether these effects are significant, for each baseline method, we performed one-sided paired t-tests to determine whether there is a significant difference between the baseline method’s error and MD-AD’s across the five test folds (**Figure 2a**). Next, in order to evaluate the contributions of each dataset to prediction performance, we performed the above procedure with different subsets of available datasets. Because ROSMAP is the only dataset with all available phenotypes, we evaluate performance specifically on ROSMAP. In **Figure 2b**, we show ROSMAP test samples’ MSE performance when trained on all subsets of ACT, MSBB, and ROSMAP training samples (following the same cross-validation procedure described above).

#### B. External dataset validation (Human)

In order to evaluate MD-AD’s ability to generalize to out of sample data, we assessed performance on three datasets: Mount Sinai Brain Bank Microarray (MSBB-M; N=1,053), Harvard Brain Tissue Resource Center (HBTRC; N=460), and Mayo Clinic Brain Bank (N=323). These datasets were collected from AMP-AD, but were left out of the original MD-AD training because they were microarray samples or lacked many neuropathology labels.

After normalizing gene expression samples from external data sets in the same way as described for the ACT, MSBB RNA Seq, and ROSMAP datasets, we then adjust the expression values to have similar distributions to our batch corrected training data sets. We evaluated the MD-AD model on our new processed data to obtain predictions for all six phenotypes. Because these three external datasets provide a sparse set of neuropathological labels, we do not have access to labels for many of the six MD-AD labels. Instead, we evaluated whether MD-AD’s predictions were consistent with the (binary) neuropathological diagnosis of AD, by aggregating MD-AD’s various neuropathology predictions into one “neuropathology score”. The “neuropathology score” was produced by first calculating percentiles across samples (within each dataset) for each neuropathological phenotype, then averaging over the six phenotypes.

**Figure 2c** shows that MD-AD provides the largest differences in neuropathology scores between individuals with and without neuropathological diagnoses of AD. We further compared neuropathology scores between AD and non-AD individuals split by age group (significance between groups shown in **Supplementary Figure 3b**)

#### C. Cross-species validation (Mouse)

To evaluate how well expression patterns predictive of neuropathology learned by MD-AD recapitulates neuropathology in mouse models. To that end, we obtained gene expression data from Matarin et al.^20^ for 30 TASTPM mice which harbor double transgenic mutation in APP and PSEN1, as well as 76 wild type mice. Data were quantile-normalized and log transformed. For this experiment, we mapped mouse to human genes (via gene symbols) for a total of 7,057 intersecting genes between our training dataset and the mouse expression data, which were again normalized to follow the same distributions as our MD-AD training data. We retrained our MD-AD model on only these 7057 genes for all MD-AD samples and then generated “neuropathology scores” for the mouse samples exactly as described in the previous section. As with out-of-sample experiments described above, we compare MD-AD to MLPs and linear models in separating neuropathology scores between TASTPM and wild type mice (Figure 2E). We also show differences in neuropathology scores between different age groups (**Figure 2d, Supplementary Figure 3c**).

#### D. Supervised embedding validation

The output of an intermediate layer of a neural network can be viewed as lower dimensional embedding of the input features. In this paper, we focus on the last shared layer of the MD-AD network because it is a supervised embedding of gene expression data which is influenced by all six training phenotypes. We evaluate the embedding compared with those generated by both singly-trained MLPs as well as unsupervised methods (i.e., K-Means and principal components analysis (PCA)) in two ways: (1) high level visualization with t-SNE, and (2) evaluating the correspondence between individual nodes and AD-related features.

##### Visualizations with t-SNE

For each of the MD-AD, MLP, and unsupervised models, we train the models on the full combined dataset. For the deep learning models, we then generate “supervised” embeddings by obtaining the output of the last shared layer (or analogous layer of the MLP model). For the unsupervised methods, K-Means and PCA, we generate an embedding of 100 dimensions to be consistent with the MD-AD and MLP models. After generating these embeddings for all samples, we then compress them to 2 dimensions via the t-SNE algorithm^21^. T-SNE Visualizations of MD-AD’s supervised embedding are shown in **Figure 3a** (left side for each phenotype), and the figure is replicated with six times, with each plot showing samples colored by neuropathological phenotype severity for each of the six phenotypes. For comparison, t-SNE visualizations for the singly-trained MLPs and unsupervised methods are shown in **Figure 3c** (colored by CERAD Score only) and colored by other phenotypes and covariates of interest in **Supplementary Figure 5**.

##### Node-phenotype correlations

To test whether MD-AD’s embedding generalizes more to AD phenotypes than the alternative methods, we compare the nodes that best capture each phenotype among MD-AD, MLPs, and unsupervised methods. We perform the following analysis with the same five training and test splits described earlier: for each of the six phenotypes used in MD-AD’s training, we identify the node in MD-AD’s last shared layer whose output is most significantly correlated with that phenotype in the training set. We then report the −log_10_(*p*-value) (after FDR correction over nodes) for the correlation between that node’s output and the training phenotype in the test set, averaged across the train/test splits. (**Figure 3a**, right side for each phenotype).

We also perform a similar analysis with higher-level AD phenotypes not used during model training: dementia diagnosis (binary variable available in all datasets), last available cognition score (controlling for age, sex, and education; only available for the ROSMAP dataset), and AD duration (i.e., time between dementia diagnosis and death; available for the ACT and ROSMAP datasets). For this analysis, we report the highest −log_10_(p-value) after FDR correction between nodes and the high-level phenotypes, average over the five test sets (**Figure 3b**).

### 4. MODEL INTERPRETATION

#### A. Constructing and annotating MD-AD consensus nodes (Figure S7)

Because deep neural networks have non-convex loss functions, randomness in our training procedure produces networks with different weights from run to run. In order to capture robust nodes and highly relevant genes, we repeat our training procedure 100 times, in order to simulate a “consensus network”. As shown in **Supplementary Figure 6a**, we construct “MD-AD consensus nodes” by clustering nodes from many runs: (1) we train 100 MD-AD networks, (2) we obtain last shared layer node outputs for all samples and normalize them (0-mean, unit variance), (3) we combine all nodes across all runs and then cluster them using k-means (where the dimensions used to calculate similarity are samples) with k=50, (4) we summarize each cluster of nodes by their medoid. Thus, for each sample, the MD-AD consensus embedding is made up of 50 nodes which are medoids of clusters generated from 100 re-trainings.

In **Supplementary Figure 4b**, we provide a visual overview of the MD-AD consensus embedding generated as described above. To provide a simple view of clusters, we select a subset of samples for which we have clear high or low pathology, excluding ambiguous cases. We include (1) individuals with Braak stage of at least 5 and CERAD scores at least 3 (i.e., “moderate”), or (2) individuals with Braak stage of 3 or lower and a CERAD score of 1 (i.e., “absent”) who are at least 85 years old and have no dementia. Case 1 captures all individuals with pathologic AD diagnoses (with and without dementia), whereas case 2 captures all individuals considered “resistant” to AD due to their old age but lack of cognitive or neurological decline (consistent with previous literature, e.g. Latimer et al. (2019)). To annotate each node in the consensus embedding, we display their correlations with various phenotypes and covariates, as well as their enrichment for REACTOME pathways.

##### Correlations

For each variable (neuropathological phenotypes, high-level AD phenotypes, and covariates), we compute the correlation −log_10_(p-value) between the variable and each consensus node output. In **Supplementary Figure 4c**, a high −log_10_(p-value) indicates that a node captures (or is highly linearly related to) a variable.

##### Pathway enrichment

Beyond relationships between nodes and phenotypes, we annotated nodes with which gene sets are relevant to their outputs. For each of the fully trained MD-AD model, we use integrated gradients (IG)^12^ to obtain sample-level gene importances for each consensus node. Note that each consensus node (as medoid within a cluster) is some node in one of the 100 re-training runs of MD-AD, thus we perform integrated gradients for the specific node in that network. By generating sample-level gene attributions for each sample, we are able to aggregate the absolute IG values across samples to obtain average gene attributions for each gene on each node. For each MD-AD consensus node, this method therefore provides us with a ranking over all genes by their importance. We then test for enrichment of REACTOME pathways^40^ in these gene rankings via gene set enrichment analysis (GSEA)^22,23^ to identify whether certain pathways seem to be involved in the activation these nodes. Enriched pathways for the MD-AD consensus nodes are shown in **Supplementary Figure 4d. Supplementary Table 2** provides detailed annotations for each node.

#### B. Identifying MD-AD’s top genes

In order to identify genes that drive MD-AD predictions, we used integrated gradients (IG)^12^ to provide importance estimates of each gene on the predicted outcomes. Again, in order to improve model stability, we calculate gene rankings based on 100 re-trainings. After each run of training, we take our trained model and apply IG for each sample to get the importance of each gene on each phenotype prediction. We then calculate a weighted average by sample (weighted by relative pathology) to compute a global importance value for each gene on each phenotype, where positive values indicate that high expression of the gene relates to more severe AD phenotypes. Finally, by averaging over all phenotypes, we obtain our final “IG score” the given round of MD-AD training. By averaging these score across 100 re-trainings, we arrive at our “consensus IG score” for MD-AD. Negative scores imply that higher expression is associated with less pathology, while positive scores imply that higher expression is associated with more pathology, according to MD-AD. We note that 100 re-trainings are more than enough to converge to a stable gene ranking (**Supplementary Figure 6c**). The top genes for MD-AD are shown in **Figure 4a**, and enriched REACTOME pathways in the top ranked MD-AD genes (via GSEA) are shown in **Figure 4b**. The full gene ranking, generated separately for each phenotype, is provided in **Supplementary Table 4**.

For comparison with a linear gene ranking method, we also calculate the correlations between each gene with each neuropathological phenotype (across all samples in our dataset), and then rank the genes by their average correlation coefficients across all six phenotypes. Comparisons between REACTOME categories represented in the top MD-AD vs correlation-based rankings are shown in **Figure 4c**.

#### C. Calculating nonlinear effects for MD-AD genes

As a deep learning method MD-AD has the capacity to identify non-linear relationships among genes’ expression levels and neuropathological phenotypes. These non-linear relationships may reveal an implicit capture of interaction effects with other covariates observable from expression data. Thus, we sought to investigate the presence of interactions between sample-level covariates and specific genes in their contributions to the MD-AD predictions. To monitor the presence of these interaction effects, we modeled the consensus IG scores as a linear combination of a gene’s expression level, a covariate of interest, and the interaction of the two. Specifically, *score_**g,i**_* = ***a*** *expr_**g,i**_* + ***b*** *feat_**i**_* + ***c*** *expr_**g,i**_ feat_**i**_* + ***d***, where *score_g,i_* is the consensus IG value for gene *g* and sample *i, expr_g,i_* is the sample *i*’s expression level for gene *g*, and *feat_i_* is sample *i*’s value for the covariate. Based on this representation, we consider there to be an interaction effect between a gene and feature on its importance in the MD-AD model if the learned ***c*** coefficient is statistically significant (*p*<.05, after FDR correction over all genes). We primarily focus on identifying an interaction effects with sex (*feat_i_* = 1 if sample *i* comes from a male), and rank interactions between genes and sex for MD-AD based on the −log10(p-value) of the interaction term.

##### Gene set enrichment

We evaluated whether sex-differential genes were enriched for the following gene sets: (1) REACTOME pathways^40^ and (2) microglial cluster gene signatures from a recent single cell RNA Seq analysis of microglial cells from autopsied aging brains^30^. To evaluate whether the list of sex-differential MD-AD genes are enriched for gene sets of interest, we use Fisher’s exact tests to evaluate the significance of overlap between all sex-differential genes and members of each gene set. Next, to evaluate whether the top MD-AD sex-differential genes are enriched for the same gene sets, we perform Fisher’s exact tests again, but this time only consider the top 100 MD-AD genes in the calculations.

### 5. BLOOD GENE EXPRESSION VALIDATION

To evaluate the ability of MD-AD to transfer to blood gene expression data, we downloaded publically available AddNeuroMed cohort data from GEO (GSE63060 and GSE63061, which we refer to as Blood1 and Blood2, respectively). Details about the AddNeuroMed samples are provided in **Supplementary Table 3**. As with the other validation datasets, each blood dataset was normalized such that each gene’s expression values have the same mean and variance as the processed MD-AD expression data. Because each blood dataset had a different set of available genes, for each dataset, we re-trained MD-AD consensus models for brain samples with only the genes available between them and blood samples (12,104 and 11,392 genes for Blood1 and Blood2 respectively). Because these blood samples came from living participants, we do not have access to the many neuropathology variables available across the brain samples. Instead, we assess whether MD-AD’s predictions align with individuals’ cognitive diagnosis of cognitively normal (CTL), mild cognitive impairment (MCI), or dementia.

We evaluate the effectiveness of the MD-AD model by comparing predicted MD-AD pathology scores between CTL and MCI individuals, and between CTL individuals and individuals with dementia via two-sided t-tests (together, and split by age). To evaluate the MD-AD embedding for blood samples, separately for each blood dataset, we obtain the last shared layer embeddings of both the MD-AD brain expression samples and blood samples from the first round of training.

## REFERENCES

1. De Jager, P. L., Yang, H. S. & Bennett, D. A. Deconstructing and targeting the genomic architecture of human neurodegeneration. Nat. Neurosci. 21, 1310–1317 (2018).

2. Gaiteri, C., Mostafavi, S., Honey, C. J., De Jager, P. L. & Bennett, D. A. Genetic variants in Alzheimer disease-molecular and brain network approaches. Nat. Rev. Neurol. 12, 413–427 (2016).

3. Marioni, R. E. et al. GWAS on family history of Alzheimer’s disease. Transl. Psychiatry 8, 0–6 (2018).

4. Jansen, I. E. et al. Genome-wide meta-analysis identifies new loci and functional pathways influencing Alzheimer’s disease risk. Nat. Genet. 51, 404–413 (2019).

5. Kunkle, B. W. et al. Genetic meta-analysis of diagnosed Alzheimer’s disease identifies new risk loci and implicates Aβ, tau, immunity and lipid processing. Nat. Genet. 51, 414–430 (2019).

6. Mostafavi, S. et al. A molecular network of the aging human brain provides insights into the pathology and cognitive decline of Alzheimer’s disease. Nat. Neurosci. 21, 811–819 (2018).

7. Zhang, B. et al. Integrated systems approach identifies genetic nodes and networks in late-onset Alzheimer’s disease. Cell 153, 707–720 (2013).

8. Mathys, H. et al. Single-cell transcriptomic analysis of Alzheimer ‘ s disease. Nature 570, 332–337 (2019).

9. Logsdon, B. A. et al. Meta-analysis of the human brain transcriptome identifies heterogeneity across human AD coexpression modules robust to sample collection and methodological approach. (2019). doi:10.7303/syn17114455

10. Blalock, E. M. et al. Incipient Alzheimer’s disease: Microarray correlation analyses reveal major transcriptional and tumor suppressor responses. Proc. Natl. Acad. Sci. U. S. A. 101, 2173–2178 (2004).

11. Katsel, P., Li, C. & Haroutunian, V. Gene expression alterations in the sphingolipid metabolism pathways during progression of dementia and Alzheimer’s disease: A shift toward ceramide accumulation at the earliest recognizable stages of Alzheimer’s disease? Neurochem. Res. 32, 845–856 (2007).

12. Sundararajan, M., Taly, A. & Yan, Q. Axiomatic attribution for deep networks. 34th Int. Conf. Mach. Learn. ICML 2017 7, 5109–5118 (2017).

13. Bennett, D. A., Schneider, J. A., Arvanitakis, Z. & Wilson, R. S. Overview and findings from the religious orders study. Curr. Alzheimer Res. 9, (2012).

14. Bennett, D. A. et al. Overview and Findings from the Rush Memory and Aging Project. Curr. Alzheimer Res. 9, 646–663 (2012).

15. Miller, J. A. et al. Neuropathological and transcriptomic characteristics of the aged brain. Elife 6, 1–26 (2017).

16. Wang, M. et al. The Mount Sinai cohort of large-scale genomic, transcriptomic and proteomic data in Alzheimer’s disease. Sci. Data 5, 1–16 (2018).

17. Johnson, W. E., Li, C. & Rabinovic, A. Adjusting batch effects in microarray expression data using empirical Bayes methods. Biostatistics 8, 118–127 (2007).

18. Mirra, S. S. et al. The consortium to establish a registry for Alzheimer’s disease (CERAD). Part II. Standardization of the neuropathologic assessment of Alzheimer’s disease. Neurology 41, 479–486 (1991).

19. Braak, H. & Braak, E. Neuropathological stageing of Alzheimer-related changes. Acta Neuropathol. 82, 239–259 (1991).

20. Matarin, M. et al. A Genome-wide gene-expression analysis and database in transgenic mice during development of amyloid or tau pathology. Cell Rep. 10, 633–644 (2015).

21. van der Maaten, L. & Hinton, G. Visualizing Data using t-SNE. J. Mach. Learn. Res. 9, 2579–2605 (2008).

22. Daly, M. J. et al. PGC-1α-responsive genes involved in oxidative phosphorylation are coordinately downregulated in human diabetes. Nat. Genet. 34, 267–273 (2003).

23. Subramanian, A. et al. Gene set enrichment analysis: A knowledge-based approach for interpreting genome-wide expression profiles. Proc. Natl. Acad. Sci. U. S. A. 102, 15545–15550 (2005).

24. Qiu, Y.-Q. KEGG Pathway Database. in Encyclopedia of Systems Biology (eds. Dubitzky, W., Wolkenhauer, O., Cho, K.-H. & Yokota, H.) 1068–1069 (Springer New York, 2013). doi:10.1007/978-1-4419-9863-7_472

25. Trabzuni, D. et al. Widespread sex differences in gene expression and splicing in the adult human brain. Nat. Commun. 4, (2013).

26. Olah, M. et al. A transcriptomic atlas of aged human microglia. Nat. Commun. 9, 1–8 (2018).

27. Chan, G. et al. CD33 modulates TREM2: Convergence of Alzheimer loci. Nat. Neurosci. 18, 1556–1558 (2015).

28. Wang, Y. et al. TREM2 lipid sensing sustains the microglial response in an Alzheimer’s disease model. Cell 160, 1061–1071 (2015).

29. Mathys, H. et al. Temporal Tracking of Microglia Activation in Neurodegeneration at Single-Cell Resolution. Cell Rep. 21, 366–380 (2017).

30. Olah, M. et al. Single cell RNA sequencing of human microglia uncovers a subset that is associated with Alzheimer’s disease. (2020).

31. Lovestone, S. et al. AddNeuroMed - The european collaboration for the discovery of novel biomarkers for alzheimer’s disease. Ann. N. Y. Acad. Sci. 1180, 36–46 (2009).

32. Johnson, E. C. B. et al. Large-scale proteomic analysis of Alzheimer’s disease brain and cerebrospinal fluid reveals early changes in energy metabolism associated with microglia and astrocyte activation. Nat. Med. 26, 769–780 (2020).

33. Lambert, J. C. et al. Genome-wide association study identifies variants at CLU and CR1 associated with Alzheimer’s disease. Nat. Genet. 41, 1094–1099 (2009).

34. Farfel, J. M. et al. Relation of genomic variants for Alzheimer disease dementia to common neuropathologies. Neurology 87, 489–496 (2016).

35. Chibnik, L. B. et al. CR1 is associated with amyloid plaque burden and age-related cognitive decline. 69, 560–569 (2011).

36. Thambisetty, M. et al. Effect of complement CR1 on brain amyloid burden during aging and its modification by APOE genotype. Biol. Psychiatry 73, 422–428 (2013).

37. Patrick, E. et al. A cortical immune network map identifies distinct microglial transcriptional programs associated with beta-amyloid and Tau pathologies. Press

38. Mirra, S. S. et al. The Consortium to Establish a Registry for Alzheimer’s Disease (CERAD): Part II. Standardization of the neuropathologic assessment of Alzheimer’s disease. Neurology 41, (1991).

39. Latimer, C. S. et al. Resistance and resilience to Alzheimer’s disease pathology are associated with reduced cortical pTau and absence of limbic-predominant age-related TDP-43 encephalopathy in a community-based cohort. Acta Neuropathol. Commun. 7, 9 (2019).

40. Jassal, B. et al. The reactome pathway knowledgebase. Nucleic Acids Res. 48, D498–D503 (2020).

