## Supplementary Figures for "Unified AI framework to uncover deep interrelationships between gene expression and Alzheimer’s disease neuropathologies"

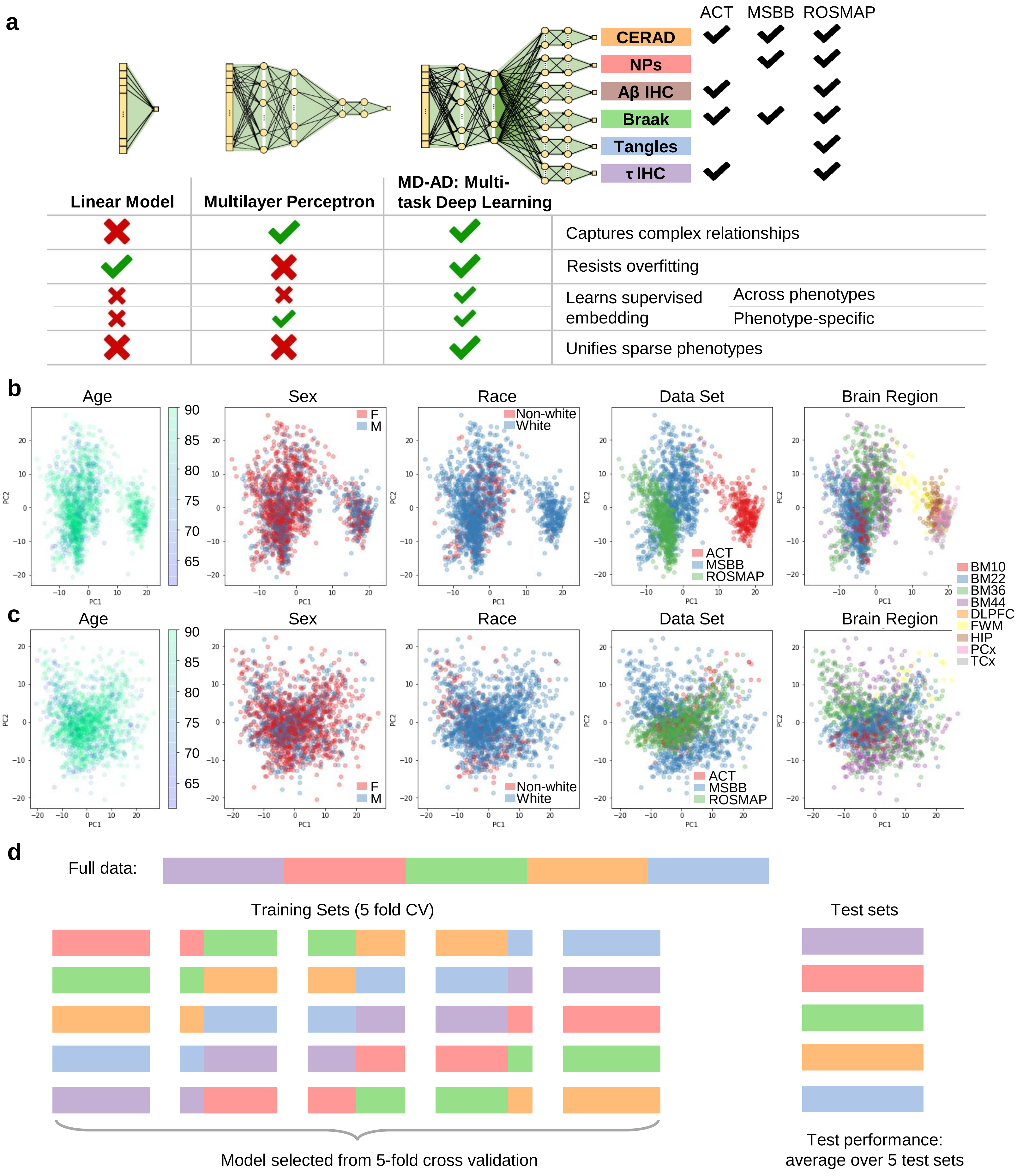


**Supplementary Figure 1. (a)** Overview of MD-AD and its advantages over traditional approaches. First two principal components of gene expression data **(a)** before and (**b**) after ComBat batch effect correction. **(c)** Overview of our cross-validation (CV) and testing scheme. We generate five separate training and test splits, and then perform cross-validation within each of the five generated training sets. For each split, we select the best hyperparameters using five-fold CV, then train the model on the training set and report the associated test performance. We then report the average over five test sets.

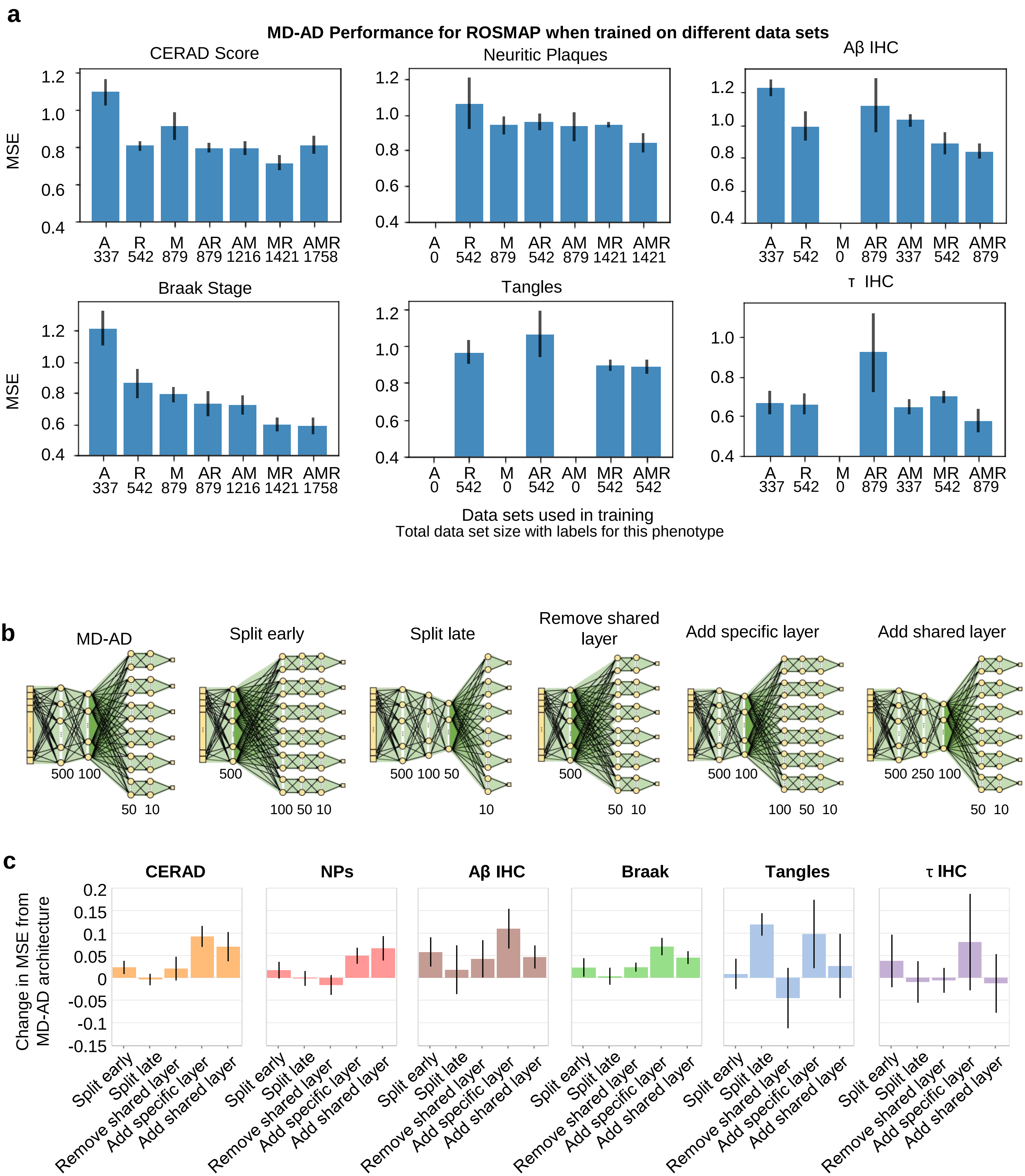


**Supplementary Figure 2. (a)** We evaluate test set performance for ROSMAP using the same training and test splits, but training restricted to different subsets of available data sets. **(b)** We experimented with several architectures for the MD-AD model. They are depicted with associated dense layer sizes. **(c)** We plot the difference in test set MSE between five alternative architectures evaluated and the final selected MD-AD architecture. Positive values indicate higher error relative to the MD-AD architecture.

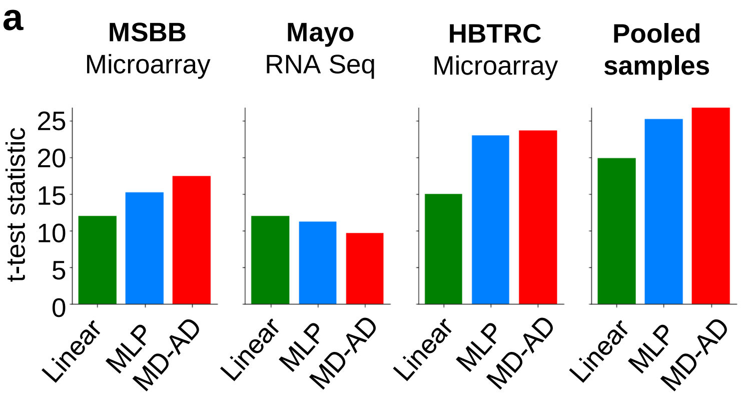


**b**

| | | <75 | | 75-85 | | >85 | |
|---|---|---|---|---|---|---|---|
| | | C | AD | C | AD | C | AD |
| <75 | C (179) | | *** | *** | *** | *** | *** |
| | AD (95) | *** | | *** | * | *** | ** |
| 75-85 | C (134) | *** | *** | | *** | | *** |
| | AD (243) | *** | * | *** | | *** | |
| >85 | C (337) | *** | *** | | *** | | *** |
| | AD (554) | *** | ** | *** | | *** | |

**c**

| | | 2 months | | | 4 months | | | 8 months | | | 18 months | | |
|---|---|---|---|---|---|---|---|---|---|---|---|---|---|
| | Group (N) | WT | HET | HO | WT | HET | HO | WT | HET | HO | WT | HET | HO |
| 2 m | WT (22) | | | | * | ** | *** | ** | *** | *** | *** | *** | *** |
| | HET (8) | | | | * | * | *** | ** | *** | *** | *** | *** | *** |
| | HO (8) | | | | | | | | | * | | * | ** |
| 4 m | WT (20) | * | * | | | | | | | ** | | ** | *** |
| | HET (8) | ** | * | | | | | | | | | | |
| | HO (8) | *** | *** | | | | | | | | | | * |
| 8 m | WT (20) | ** | ** | | | | | | | * | | * | ** |
| | HET (8) | *** | *** | | | | | | | | | | *** |
| | HO (8) | *** | *** | * | ** | | | * | | | | | |
| 18 m | WT (14) | *** | *** | | | | | | | | | | * |
| | HET (8) | *** | *** | * | ** | | | * | | | | | |
| | HO (6) | *** | *** | ** | *** | | * | ** | *** | | * | | |

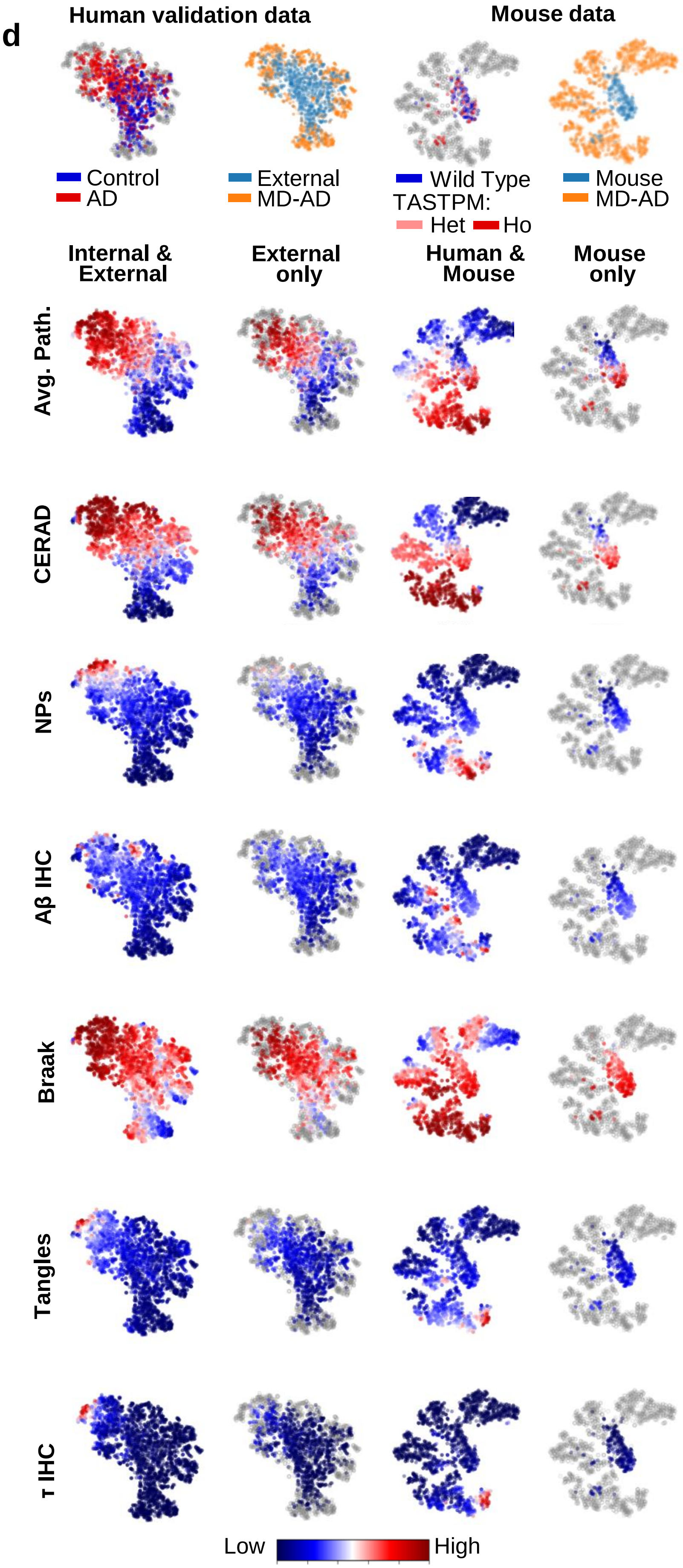


**Supplementary Figure 3.** **(a)** *t*-test statistics measuring differences between each model's predicted neuropathology scores for AD-diagnosed vs. control individuals. **(b)** Significance of *t*-test statistics measuring between-group differences as shown in boxplots in **Figure 2c** and **(c) Figure 2d** (*: $p$<.05, **: $p$<.01, ***: $p$<.001). **(d)** t-SNE plots of embedded samples for external and mouse data sets.

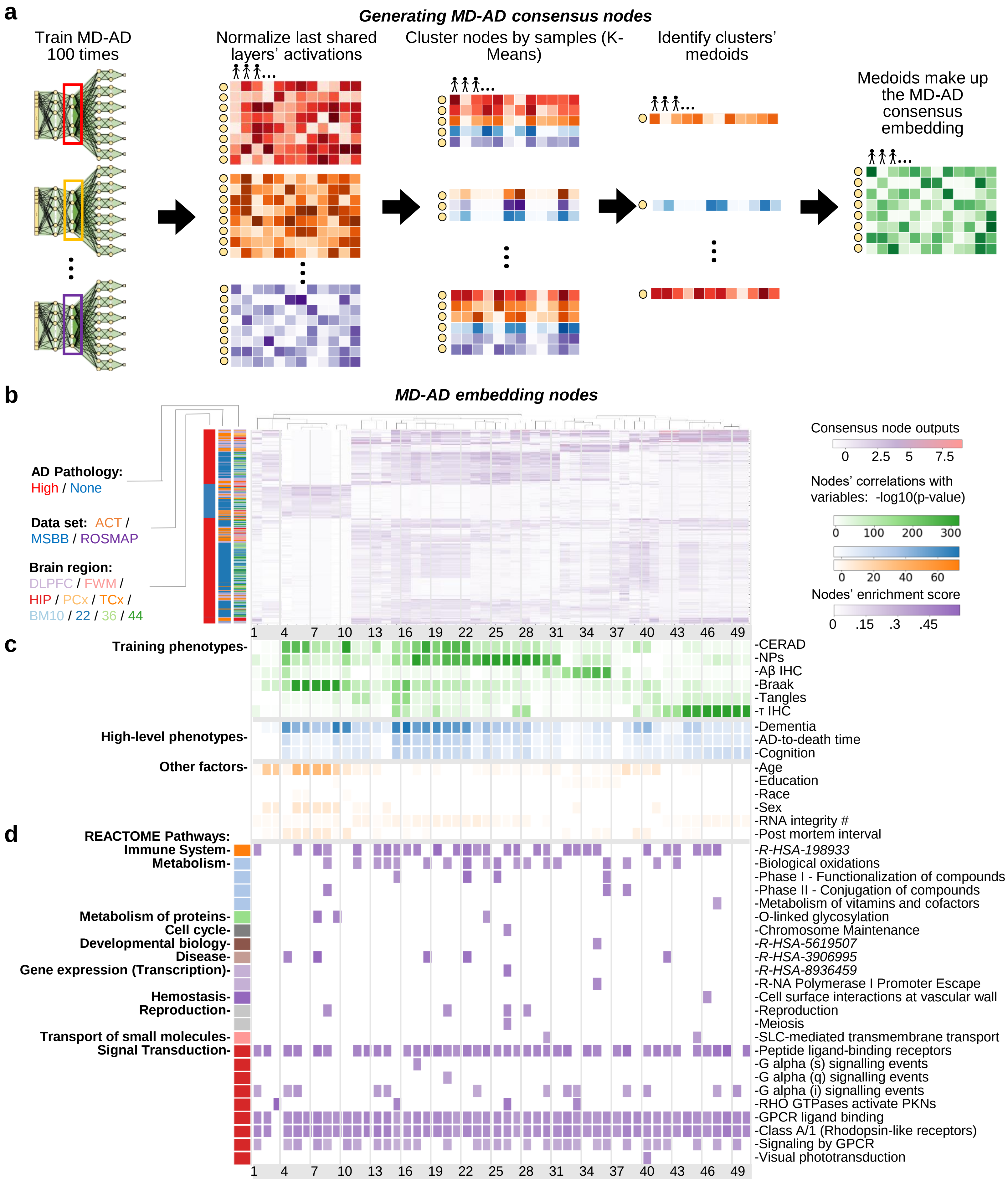


**Supplementary Figure 4.** Generating and annotating MD-AD "consensus" nodes. **(a)** Illustration of how we generate MD-AD consensus nodes. **(b)** Bi-clustered last shared layer consensus node embeddings. **(c)** Correlations between consensus nodes and phenotypes of interest. **(d)** Nodes' GSEA enrichment score (ES) for REACTOME pathways: For each node, we obtain integrated gradients weights from each gene. We display only cells with |ES|>.2 and *p*<.05 after FDR correction (across nodes). REACTOME pathways with long names are indicated by their REACTOME stable IDs.

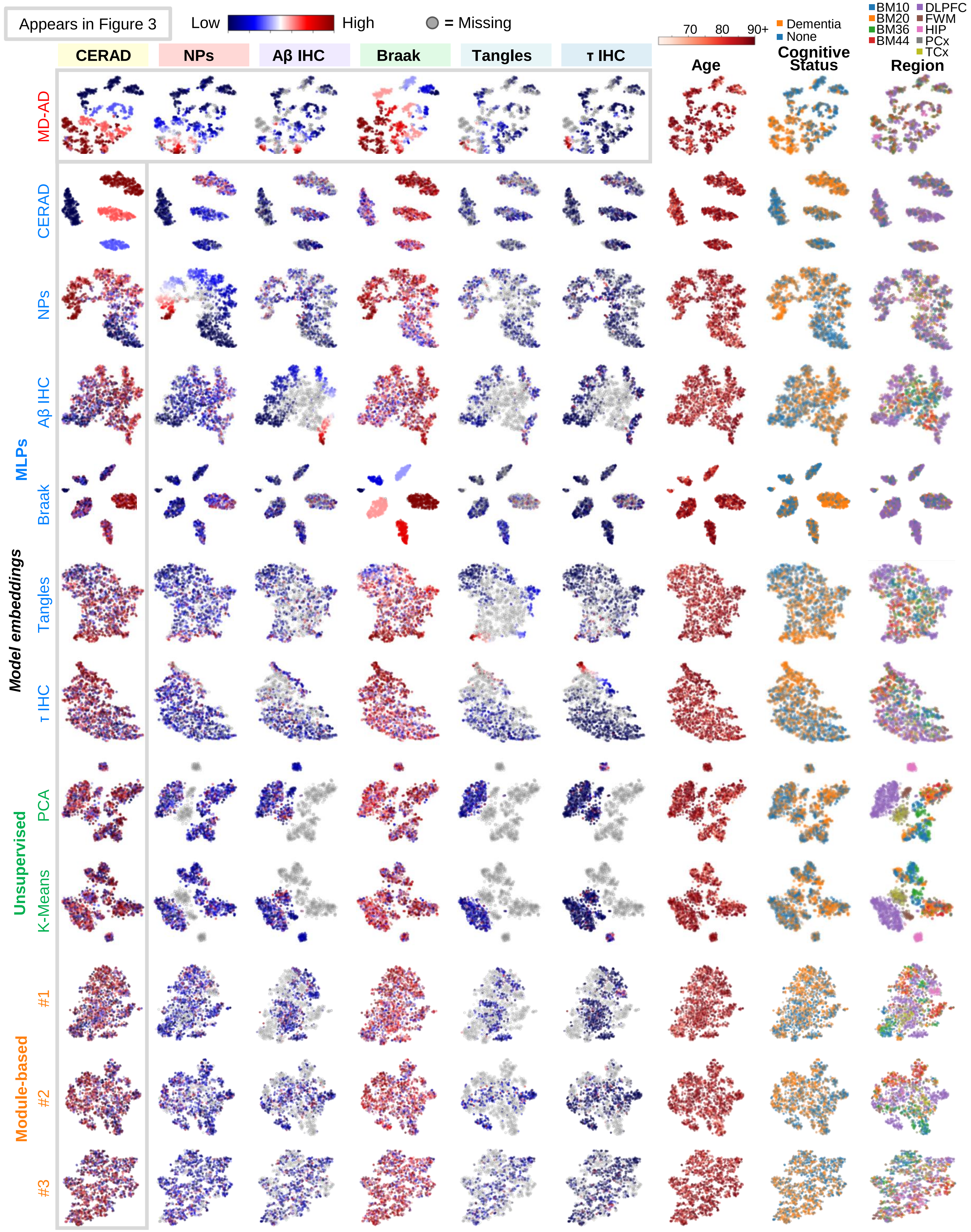


**Supplementary Figure 5.** *t*-SNE representation of all methods' embeddings colored by pathology values

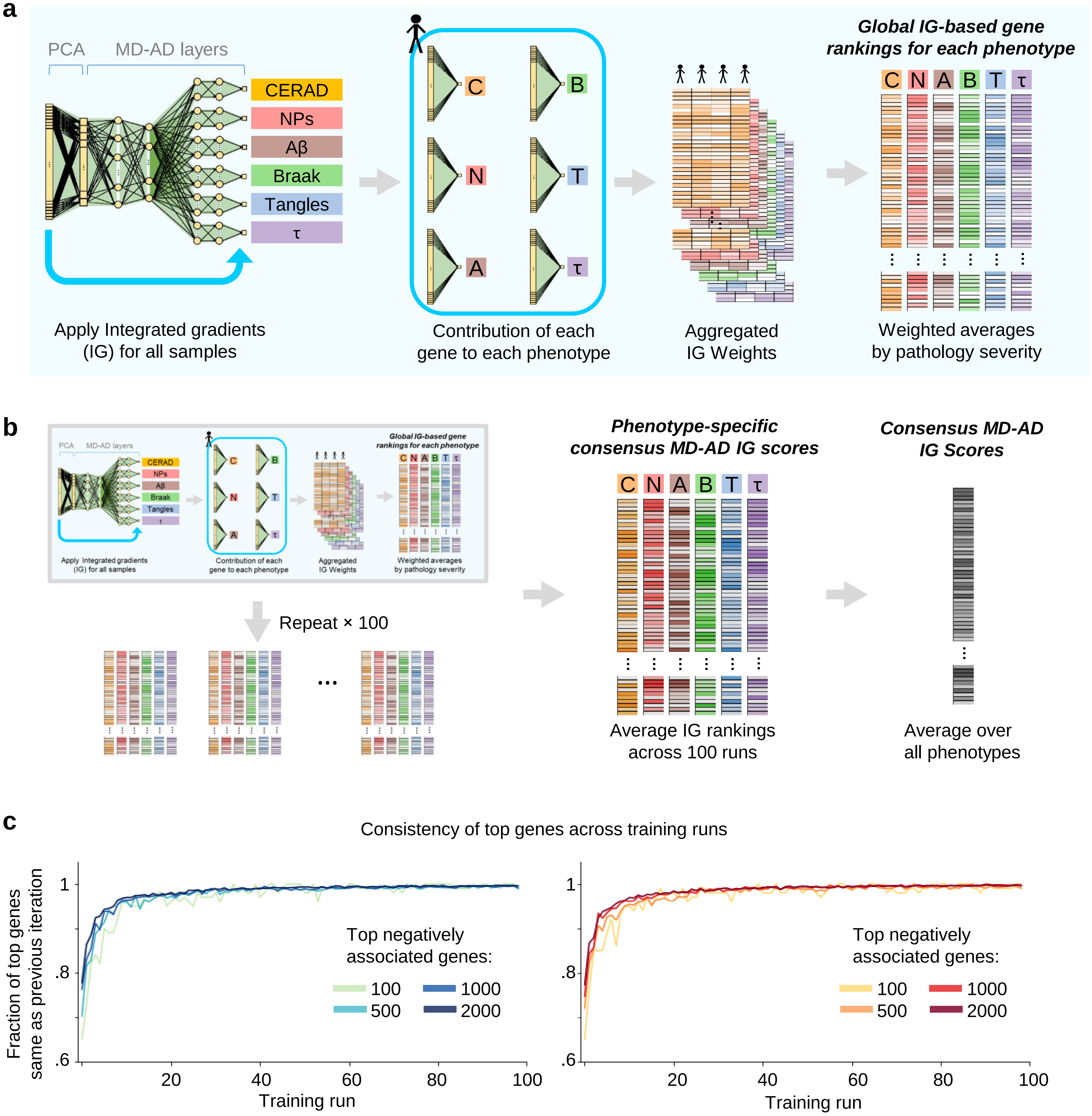


**Supplementary Figure 6.** Illustration of how MD-AD "consensus" gene scores are generated. **(a)** For a single training run, we aggregate IG scores across samples to obtain a ranking over genes for each phenotype. **(b)** We aggregate IG gene scores across 100 re-trainings of MD-AD. **(c)** Consistency of top genes from aggregating IG gene scores across multiple training runs.

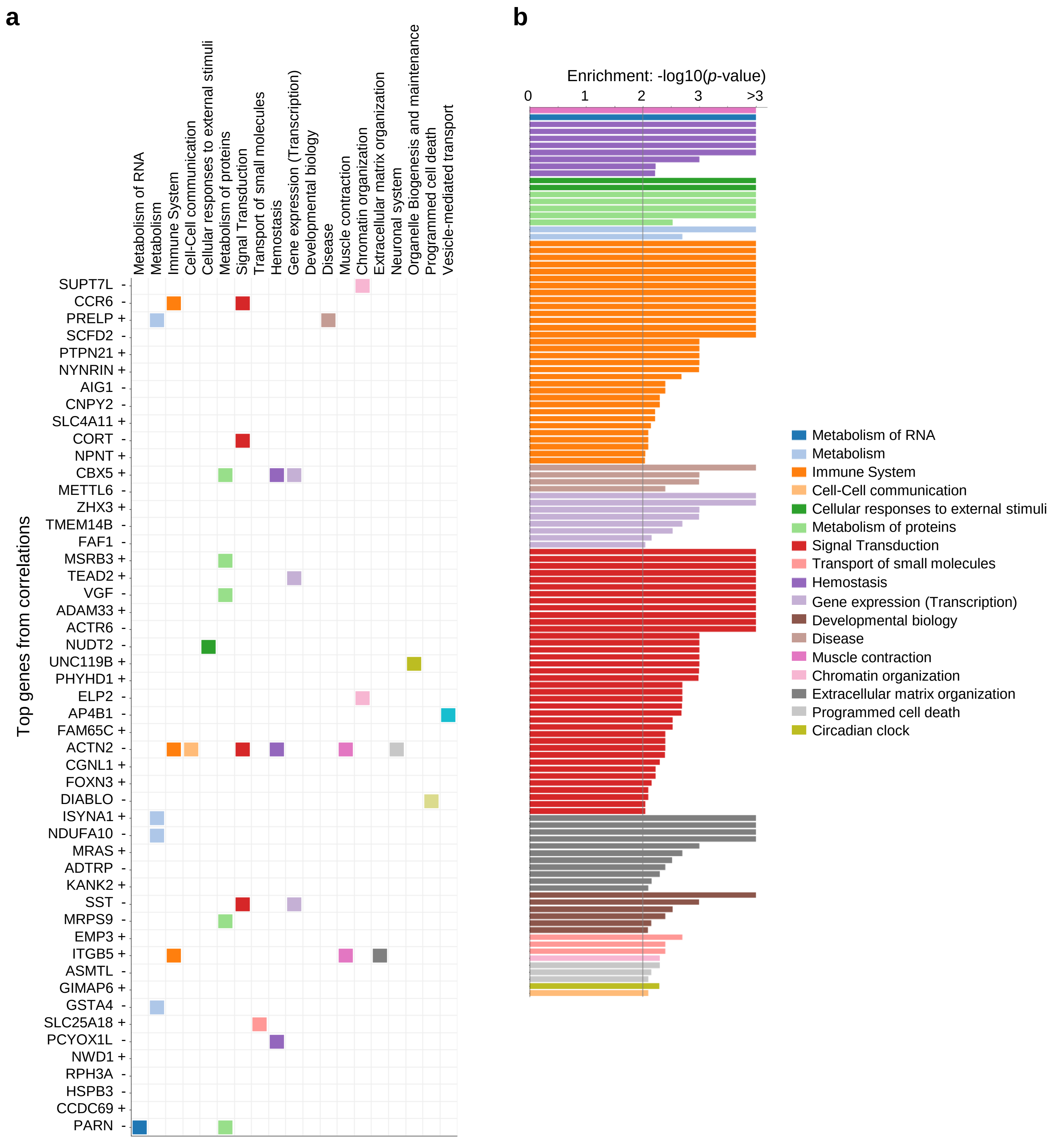


**Supplementary Figure 7**. Top genes and REACTOME pathway enrichment for correlation-based ranking: (**a**) Top 50 genes ranked by correlations between expression and pathology. (**b**) GSEA enrichment *p*-values for pathways enriched in correlation-based ranking.

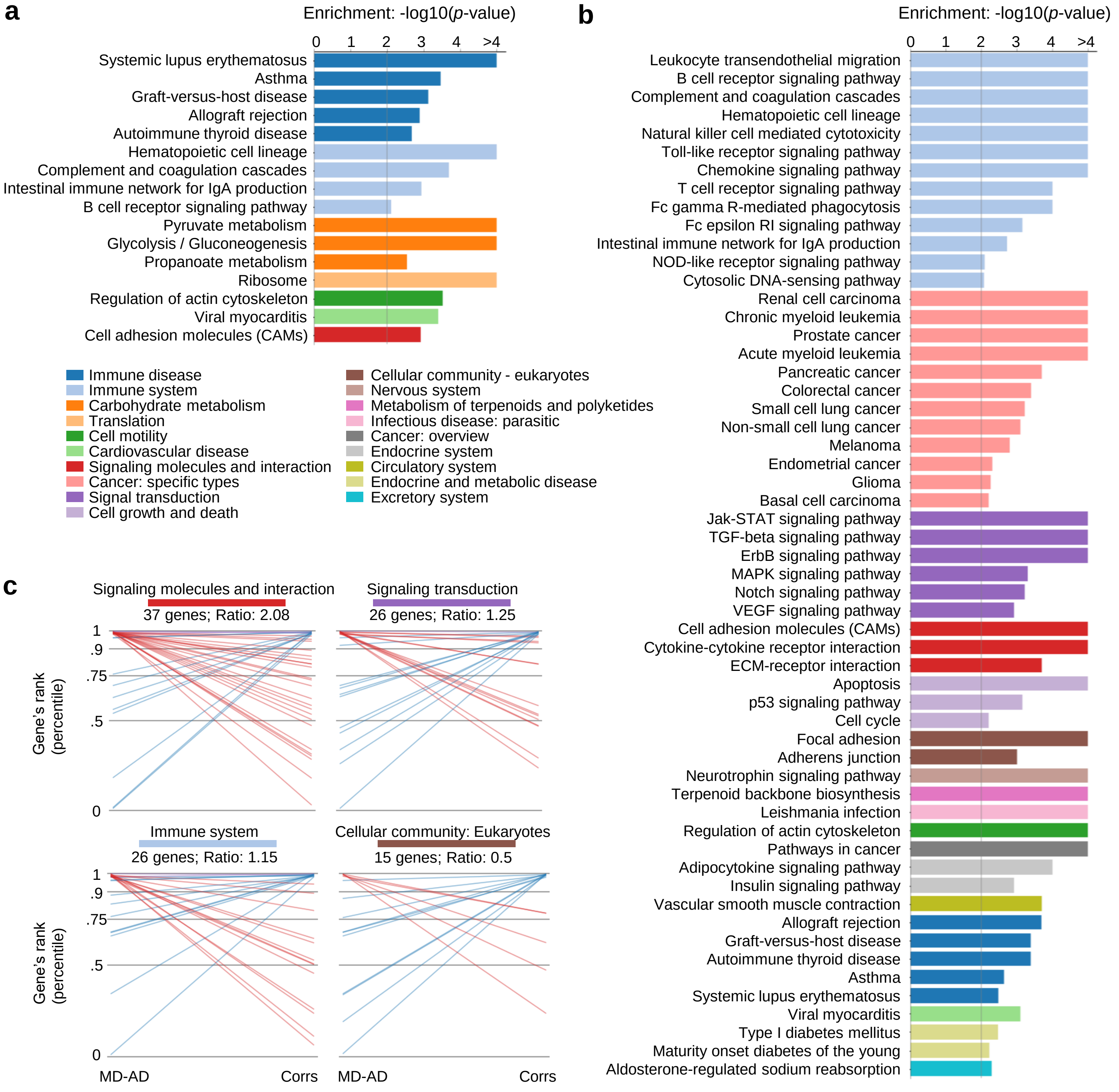


**Supplementary Figure 8**. (**a**) GSEA enrichment *p*-values for **KEGG** pathways enriched in MD-AD gene ranking, (**b**) GSEA enrichment *p*-values for pathways enriched in correlation-based ranking, (**c**) Comparison of top genes for MD-AD vs correlations. For MD-AD and correlation-based rankings, we identify all genes in the top 2% of the ranking, and then check their membership in KEGG categories.

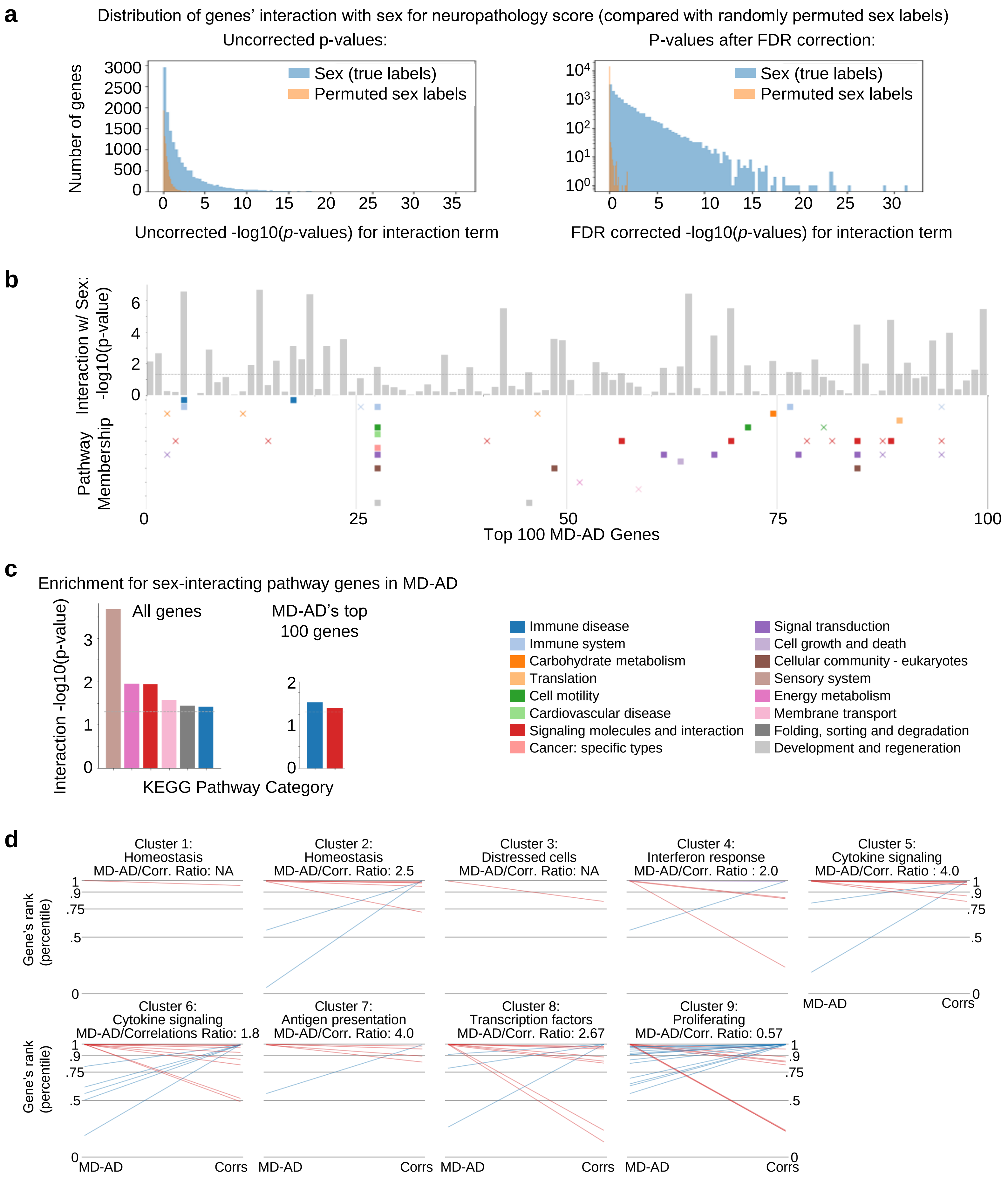


**Supplementary Figure 9.** Additional details for sex interaction results. **(a)** Distribution of genes' interactions with sex for MD-AD scores. For each gene, we compute the –log10(p-value) for the interaction term between the gene's expression and sex. For comparison, we show the distribution from an experiment with all sex labels shuffled across samples. **(b)** Replicated results from **Figure 5a** with KEGG categories. **(c)** Replicated results from **Figure 5b** with KEGG categories. **(d)** Comparison of top 1% ranked genes from MD-AD vs a correlation-based approach for microglial cluster members.

a

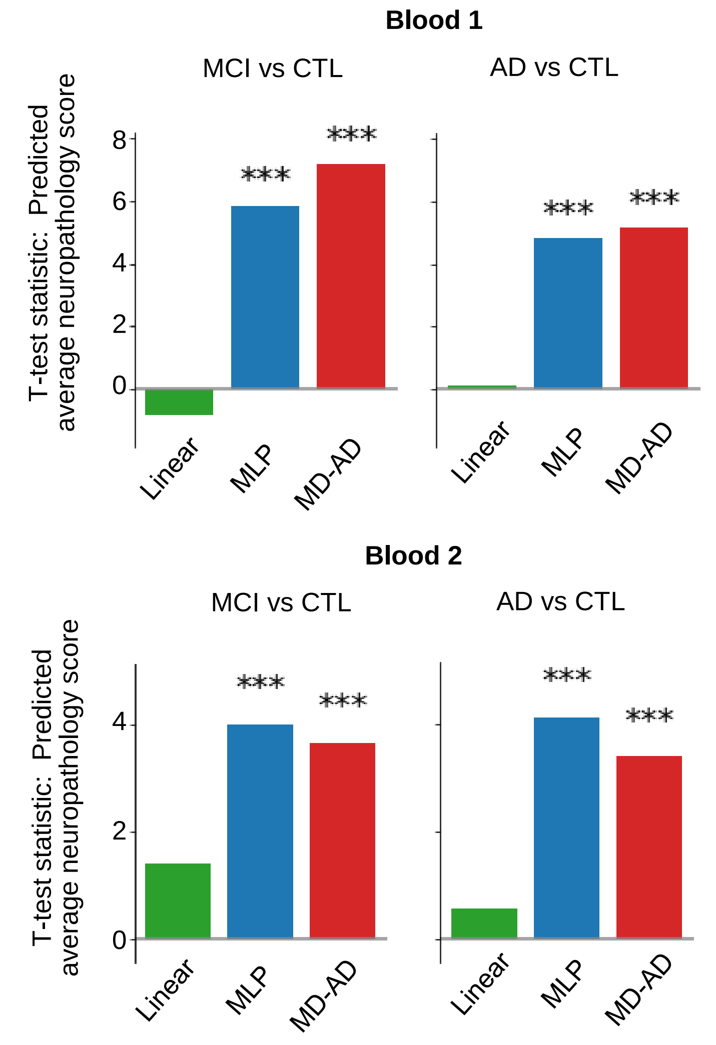


b

| | | <74 | | | [74-80) | | | 80+ | | |
|---|---|---|---|---|---|---|---|---|---|---|
| | Group (N) | CTL | MCI | AD | CTL | MCI | AD | CTL | MCI | AD |
| <74 | CTL (113) | | *** | *** | | *** | *** | | *** | *** |
| | MCI (64) | *** | | | *** | | * | ** | | |
| | AD (89) | *** | | | *** | | | * | | |
| [74-80) | CTL (82) | | *** | *** | | *** | ** | | ** | ** |
| | MCI (63) | *** | | | *** | | | ** | | |
| | AD (90) | *** | * | | ** | | | | | |
| 80+ | CTL (43) | | ** | * | | ** | | | | |
| | MCI (62) | *** | | | ** | | | | | |
| | AD (105) | *** | | | ** | | | | | |

c

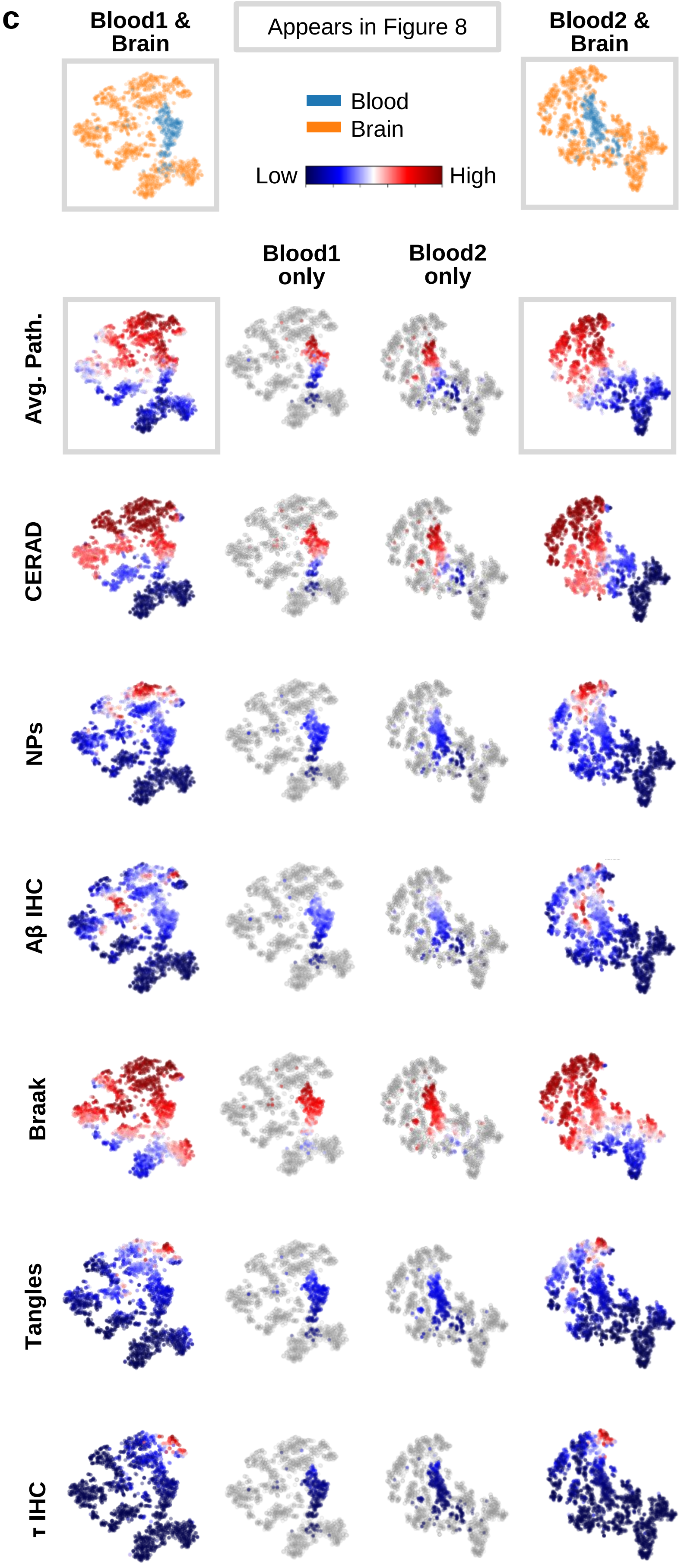


**Supplementary Figure 10**. (**a**) t-test statistics for comparisons among MD-AD predicted neuropathology and cognitive states, separately for blood datasets. (**b**) pair-wise *t*-test *p*-values from **Figure 7b**. (*: $p<.05$, **: $p<.01$, ***: $p<.001$) **(c)** Embeddings from MD-AD's last shared layer for brain and blood data. Each plot is colored by dataset or predicted pathology severity for various phenotypes. Pathology severity plots are produced with and without MD-AD training (brain) samples.
